# TR(acking) individuals down: exploring the effect of temporal resolution in resting-state functional MRI fingerprinting

**DOI:** 10.1101/2023.11.20.565789

**Authors:** Barbara Cassone, Francesca Saviola, Stefano Tambalo, Enrico Amico, Silvio Sarubbo, Dimitri Van De Ville, Jorge Jovicich

## Abstract

Functional brain fingerprinting has emerged as an influential tool to quantify reliability in neuroimaging studies and to identify cognitive biomarkers in both healthy and clinical populations. Recent studies have revealed that brain fingerprints reside in the timescale-specific functional connectivity of particular brain regions. However, the impact of the acquisition’s temporal resolution on fingerprinting remains unclear. In this study, we examine for the first time the reliability of functional fingerprinting derived from resting-state functional MRI (rs-fMRI) with different whole-brain temporal resolutions (TR = 0.5, 0.7, 1, 2, and 3 s) in a cohort of 20 healthy volunteers. Our findings indicate that subject identifiability within a fixed TR is successful across different temporal resolutions, with the highest identifiability observed at TR 0.5 and 3 s. We discuss this observation in terms of protocol-specific effects of physiological noise aliasing. We further show that, irrespective of TR, associative brain areas make substantial contributions to subject identifiability, whereas sensory-motor regions become influential only when integrating data from different TRs. We conclude that functional connectivity fingerprinting derived from rs-fMRI holds significant potential for multicentric studies also employing protocols with different temporal resolutions. However, it remains crucial to consider fMRI signal’s sampling rate differences in subject identifiability between data samples, in order to improve reliability and generalizability of both whole-brain and specific functional networks’ results. These findings contribute to a better understanding of the practical application of functional connectivity fingerprinting, and its implications for future neuroimaging research.

## 1. Introduction

In recent years, brain connectome fingerprinting has shown increased promise as a tool for understanding underlying mechanisms of the human brain, in particular those related to functional connectivity (FC), which represents the statistical interdependency between brain regions’ activity across time (Friston, 1994). In their seminal work, Finn and colleagues (2015) were among the first to show that an individual’s FC profile is not only unique, but also reliable enough to be identified and distinguished with high accuracy from a larger sample of similar healthy individuals. Following studies expanded on the topic by showing that functional patterns are useful to detect subject-level biomarkers of behavioral outcome and cognitive performance (Amico and Goñi, 2018; Romano et al., 2022; Sorrentino et al., 2021; Svaldi et al., 2021; Troisi Lopez et al., 2022), opening up the possibility to predict clinical scores and contributing to personalized treatments planning (Castellanos et al., 2013; Fernandes et al., 2017; Smith et al., 2015) and biometric systems (Fraschini et al., 2015; Rocca et al., 2014).

Another relevant application of the study of brain fingerprints relates to one essential aspect and source of concern for numerous domains of scientific research (Begley & Ioannidis, 2015), that is reproducibility and generalizability of results. In particular, Amico and Goñi (2018) introduced the “identifiability framework” for assessing and increasing subject identification across visits, by running principal component analysis on a group’s functional connectomes, and removing “noise” components while retaining only components necessary for subject identification. This procedure has been shown to increase test-retest reliability across different tasks (Amico and Goñi, 2018; Rajapandian et al., 2020), scanning lengths (Amico and Goñi, 2018), MRI scanners and sites (Bari et al., 2019), network properties (Rajapandian et al., 2020), imaging modalities, namely fMRI and MEG, FC measures and frequency bands (Sareen et al., 2021). Moreover, specific brain regions have been found to give a different contribution to fingerprinting (Amico and Goñi, 2018; Romano et al., 2022; Sorrentino et al., 2021; Svaldi et al., 2021; Troisi Lopez et al., 2022; Van de Ville et al., 2021), which is maximized at specific timescales, consistently with intrinsic neural firing patterns (Gao et al., 2020) and underlying mental processes (Van de Ville et al., 2021). This raises the question of how functional acquisition timescales may affect subject identification within this functional fingerprinting framework.

Understanding the effect of temporal resolution on fingerprinting of resting-state fMRI (rs-fMRI) holds relevance for several reasons. Firstly, rs-fMRI has gained widespread use in both basic and clinical research, making it crucial to comprehend the factors influencing its outcomes (Lee et al., 2013a; Smitha et al., 2017). Secondly, the growing influence of the open-science policy has encouraged access to multicentric rs-fMRI datasets acquired using different temporal resolutions, emphasizing the need to examine the implications of this factor on data interpretation and integration (for example: https://openfmri.org, https://adni.loni.usc.edu, http://www.humanconnectomeproject.org, http://www.developingconnectome.org, https://www.ukbiobank.ac.uk, https://fcon_1000.projects.nitrc.org/indi/CoRR/html/ among various other public rs-fMRI data sources). Lastly, advancements in rapid parallel imaging technologies (Larkman et al., 2001) have enabled neuroscientists to obtain whole-brain images with sub-second sampling rates, thus allowing the investigation of neural dynamics with greater precision and flexibility (Akin et al., 2017; Lee et al., 2013b; Zalesky et al., 2014). However, the reliability of fast fMRI remains unclear. On the one hand, acquiring a higher number of timepoints without increasing the scanning duration enhances statistical power (Dowdle et al., 2021; Feinberg et al., 2010; Posse et al., 2012; Smith et al., 2013). On the other hand, speed comes at the cost of lower signal-to-noise ratio per time frame (Barth et al., 2016; Boubela et al., 2014; Edelstein et al., 1986; Feinberg & Setsompop, 2013; Preibisch et al., 2015) with respect to conventional fMRI protocols (TR ∼2-3 s), with the ultimate result of decreasing the statistical validity of inferences about intrinsic functional connectivity (Corbin et al., 2018). To the best of our knowledge, very few studies have assessed test-retest reliability in fast fMRI (Jahanian et al., 2019) or in comparisons between different TRs. In the few instances where such comparisons were made, data were downsampled (Birn et al., 2013; Houtari et al., 2019; Shah et al., 2016) rather than acquired in distinct fMRI sessions with different temporal resolutions, which is not equivalent. Consequently, the impact of acquisition’s temporal resolution on rs-fMRI fingerprinting remains largely unexplored.

The relevance of this research question lies not only in the potential implications for the reproducibility and generalizability of fMRI results, but it is also linked to the open field of investigation of the temporal aspects of brain fingerprints. In general, fast fMRI has been proven to better capture complex temporal features of the fMRI signal which carry meaningful information about brain states (Dowdle et al., 2021; Yang & Lewis, 2021). Interestingly, in the specific framework of connectome fingerprinting, fast fMRI has provided insights into which specific time scales of connectivity patterns are more highly related to the individuality of cognitive functions (Van de Ville et al., 2021). Evaluating how brain fingerprints are affected by temporal resolution may shed light on what exactly is the information encoded in brain connectomes that ultimately makes us unique, and, specifically, weather this information unfolds in possibly preferential windows in time, when one individual’s brain is maximally identifiable (Van de Ville et al., 2021). In this scenario, exploring which temporal resolution can optimize the investigation of brain fingerprints’ time scales, in a network- and region-specific fashion, could advance our knowledge on mental process and cognition, with potential applications in the emerging fields of precision medicine.

In this study, we extend the investigation of temporal features of spontaneous brain fMRI fingerprints by addressing the question of whether the temporal resolution (TR) of rs-fMRI acquisitions’ influences healthy subject identification. We thus present the application of the “identifiability framework” (Amico and Goñi, 2018) to a nowadays still unexplored scenario. Specifically, for each subject (N = 20), we collected rs-fMRI data with five different TRs, ranging from 0.5 to 3 s. Our analyses focused on evaluating the test-retest reliability at the whole-brain level by considering the following aspects: i) the impact of TR on subject identifiability when comparing sessions acquired with the same TR; ii) the influence of TR on subject identifiability when comparing acquisitions with different TRs; iii) the effect of the number of acquired timepoints (i.e., volumes) on test-retest reliability; iv) the differential contribution of specific brain regions and functional networks to subject identifiability, regardless of TR, and to TR identifiability across subjects.

## 2. Materials and methods

A subset of the data used in the current study was also employed in a previous work (Saviola et al., 2022), available on bioRxiv. Therefore, participants’ demographics, data acquisition, and pre-processing steps were already partially described there but are reiterated here for the sake of completeness and with the necessary modifications.

### 2.1 Participants

A total of twenty healthy volunteers (10 females, age: 24 ± 3 years, 4 left-handers) without neurological and/or psychiatric disease history gave written informed consent to participate in this study, which was approved by the Ethical Committee of the University of Trento, Italy.

### 2.2 Magnetic resonance imaging acquisition

Neuroimaging data were acquired with a 3 Tesla Siemens Magnetom Prisma (Siemens Healthcare, Erlangen, Germany) whole body MRI scanner equipped with a 64-channel receive-only head-neck RF coil. Each participant underwent one structural T1-weighted standard MPRAGE (TR/TE = 2.31 s/3.48 ms, 1 mm isotropic voxels), and five acquisitions of resting-state functional MRI (rs-fMRI, TE = 28 ms, 3 mm isotropic voxels, FA = 59 degree, simultaneous multi slice acceleration factor = 6, 56 slices, fat suppression, interleaved slice acquisition, 0 mm slice gap, anterior-to-posterior phase-encoding) during the same session. Acquisition parameters for the five rs-fMRI runs were identical except for TR, which was manipulated such that the total scanning time was kept constant (7.4 minutes), thus affecting the total number of brain functional volumes (NoV acquired) per acquisition: TR(s)/NoV = 0.5/905; 0.7/646; 1/452; 2/226; 3/150. A phase and magnitude double-echo gradient echo structural sequence was used to derive a magnetic field map for geometric distortion correction (TR = 682 ms, TE_1_ = 4.2 ms, TE_2_ = 7.4 ms, 3 mm isotropic voxels).

### 2.3 Data preprocessing and functional connectivity matrices

Brain rs-fMRI data were preprocessed using FSL (Jenkinson et al., 2012) and following standard steps: i) slice timing and head motion correction; ii) T1-weighted image tissue segmentation; iii) co-registration of the rs-fMRI time-series to the T1-weighted image; iv) rs-fMRI temporal band-pass filtering ([0.01-0.3] Hz); v) nuisance regression of the 6 head motion parameters, white matter, and cerebrospinal fluid signals; vi) normalization to standard MNI template space; vii) spatial smoothing with a 6 mm FWHM kernel size.

After pre-processing, rs-fMRI images were parcellated using the Glasser multimodal parcellation atlas (Glasser et al., 2016), consisting of 360 cortical parcels. For full gray matter coverage (Amico & Goñi, 2018), we added 19 parcels from a subcortical atlas provided by the Human Connectome Project (HCP; Van Essen et al., 2013, release Q3, filename “Atlas_ROI2.nii.gz”).

FC matrices were calculated for each subject and TR by estimating the Pearson’s correlation coefficients between all parcels’ rs-fMRI time courses. The FC matrices were symmetric and were kept neither thresholded nor binarized in the following analysis (Amico & Goñi, 2018).

### 2.4 Group-Level Principal Component Analysis

In order to investigate the effect of temporal resolution on individual functional connectivity identifiability, we performed two types of fingerprinting analysis (**Figure 1**): i) a <u>within-TR analysis</u> considering, for every subject and every TR separately, FC matrices computed on the first half of the rs-fMRI time course (half volumes) as test, and FC matrices computed on the second half of the rs-fMRI time course as retest (Amico & Goñi, 2018); ii) a <u>between-TR analysis</u> considering, for every subject and all TR pairwise comparisons (i.e., TR 0.5 versus TR 0.7, TR 0.5 versus TR 1, etc.), FC matrices computed on one TR data as test, and FC matrices computed on the other TR data as retest. Note that since labeling a set of FC matrices as test or retest in a specific comparison did not elicited differences in analysis and results, we here report findings only from one possible test-retest combination (e.g., only TR 0.5 s versus TR 0.7 s results are reported because they are identical to TR 0.7 s versus TR 0.5 s results).

Since the rs-fMRI runs were identical in the total scanning time but they differed in TR, FC matrices were estimated on a different number of brain volumes, which could affect subject identifiability (Anderson et al., 2011; Birn et al., 2013; Noble et al., 2017, Van de Ville et al., 2021). To evaluate this, we repeated both the within-TR and the between-TR analysis using only the first 150 volumes (i.e. the number of volumes of the slowest TR) of each TR time course. This disentangled the effect of number of volumes from the effect of temporal resolution on individual FC identifiability.

**Figure 1.**
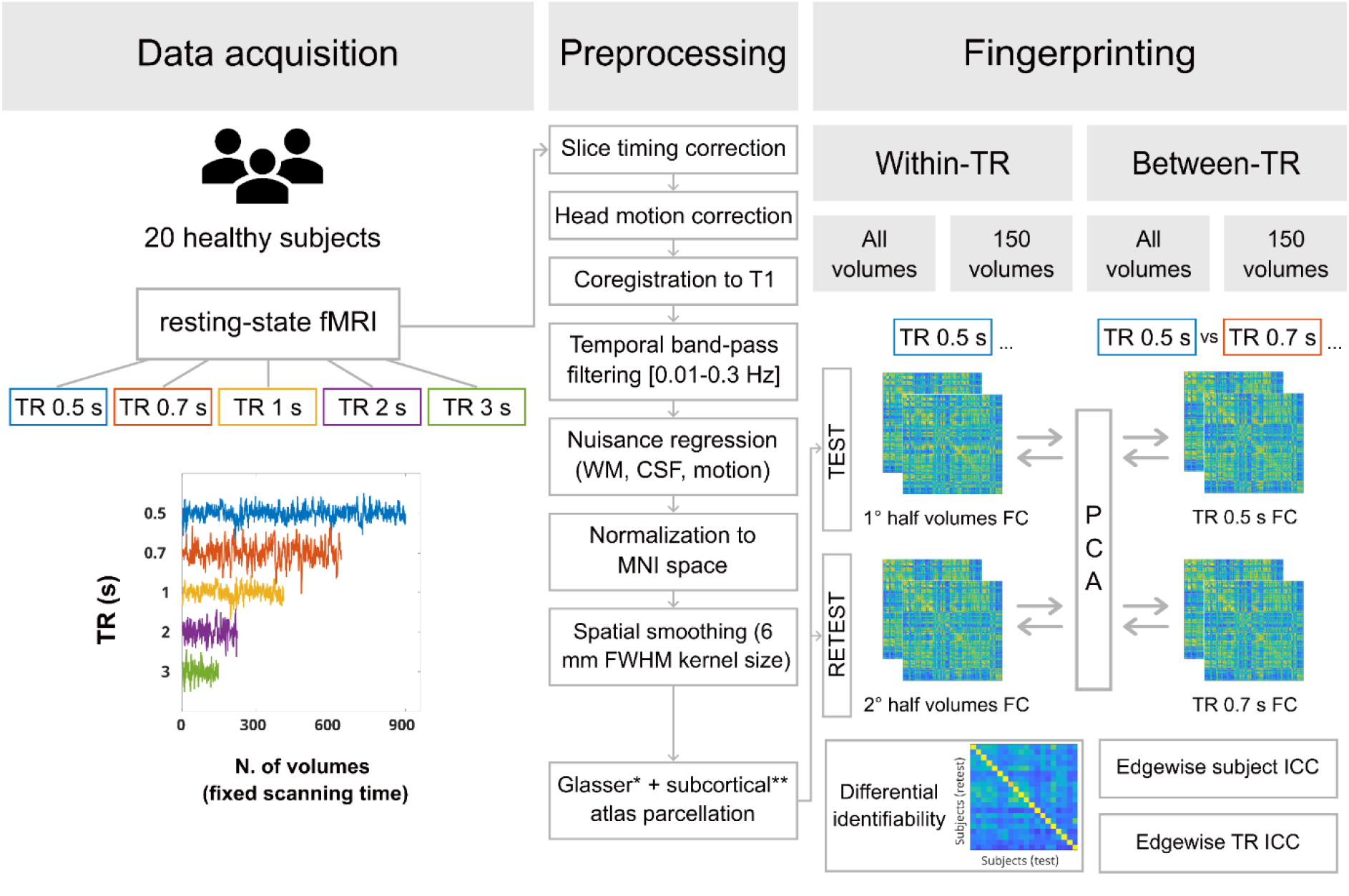
Analysis workflow. Resting-state fMRI data were acquired for each subject during five acquisitions differing in TR (TR = 0.5 s, 905 volumes; TR = 0.7 s, 646 volumes; TR = 1 s, 452 volumes; TR = 2 s, 226 volumes; TR = 3 s, 150 volumes). After preprocessing, parcellation was performed on the basis of an atlas combining Glasser parcellation (Glasser et al., 2016) and a subcortical parcellation provided by the Human Connectome Project. Two fingerprinting analyses were performed. Within-TR analysis: after parcellation, for each TR separately, time course was split in two halves, and two sets of functional connectivity (FC) matrices (test and retest) were computed as input for the group-level Principal Component Analysis (PCA). FC matrices were then reconstructed using the optimal number of PCA components. Between-TR analysis: after parcellation, for each TR pairwise comparison, the whole time course from two different TR runs (test and retest) was used to compute FC matrices. The optimal number of components resulting from the group-level PCA were used to reconstruct back each FC matrix. For both the within-TR and the between-TR analysis, Pearson’s correlation coefficients were computed between test and retest sets of FC matrices, in order to create an identifiability matrix for each TR condition and TR pairwise comparison. Finally, edgewise Intraclass Correlation (ICC) was computed in two different ways, resulting in a subject and a TR ICC. The whole fingerprinting analysis was repeated using all volumes and only the first 150 volumes of each TR session.

For both the within- and between-TR analysis, when using both all volumes and only the first 150 volumes, we applied the denoising procedure for maximizing connectivity fingerprints in human connectomes first introduced by Amico and Goñi (2018). For each subject, two FC matrices (test and retest as described in the previous paragraphs) were taken as input, their upper triangle was vectorized, and then added to a matrix having sessions (double the sample size) as columns and FC edge weights as rows. A Principal Component Analysis (PCA) algorithm (Jolliffe, 2014) was then applied to extract the optimal number of connectivity-based components to maximize subject identifiability, which were then used to reconstruct back the FC matrices of each subject.

Except for data preprocessing, analysis was conducted using MATLAB 2017b. The code for performing the fingerprinting analysis was adapted from the scripts publicly available at Enrico Amico’s GitHub repository (https://github.com/eamico). For privacy reasons, the dataset used for the current study cannot be shared. Upon acceptance, code will be make publicly available through GitHub (https://github.com/CIMeC-MRI-Lab).

### 2.5 Whole-brain connectome fingerprinting: differential identifiability and success rate

In order to assess the identifiability of FC profiles of single individuals among the entire sample at the whole-brain level, an identifiability matrix (Amico & Goñi, 2018) was created where rows referred to subjects’ FCs in the test session, and columns to subjects’ FCs in the retest session, and the Pearson’s correlation coefficient is used to compute their similarity. The average of the main diagonal elements of the identifiability matrix is defined as “self-identifiability” or “*Iself*” since it is a measure of the similarity of the same subject’s FC profile between test-retest data. The average of the main off-diagonal elements of the identifiability matrix, instead, is defined as “*Iothers*” and represents the similarity of FC profiles between test-retest data across subjects. The assumption of the functional connectome fingerprint is that FC should be, overall, more similar between test-retest sessions of the same subject than between different subjects. Thus, a “differential identifiability” or “*Idiff*” measure (*Idiff* = (*Iself* – *Iothers*) * 100) has been proposed as a group estimate of how much an individual FC profile is identifiable amongst the whole sample (Amico & Goñi, 2018). With this formalism, *Idiff* represents a continuous score for the level of individual whole-brain fingerprinting present on a set of test-retest functional connectomes. We also computed a binary identification score, namely “success rate” (*SR*), which is defined as the percentage of subjects whose identity was correctly predicted out of the total number of subjects (Finn et al., 2015). Consistent findings between *Idiff* and *SR* would suggest the possibility to generalize our subject identification results to different identifiability scores (Sareen et al., 2021).

### 2.6 Local contributions to fingerprinting: the role of individual edges and networks

Besides whole-brain FC fingerprinting, we also investigated if identifiability was affected by specific connectome edges (functional correlations between brain parcellation pairs) or networks (grouping edges) to subject identifiability. In particular, we wanted to see if particular edges or networks boosted the identifiability between subjects (regardless of TR), or the identifiability across TRs (regardless of subjects).

To do this, we computed the intraclass correlation coefficient (ICC; Bartko, 1966; McGraw & Wong, 1996), as in previous studies (Amico & Goñi, 2018; Bari et al., 2019; Sareen et al., 2021; Van de Ville et al., 2021). ICC is a statistical measure of the agreement between units of (or ratings/scores) of different groups (or raters/judges). The stronger the agreement, the higher its ICC value.

We employed ICC to quantify the extent to which individual edges can separate between subjects (units) regardless of the TRs (raters). Specifically, for each edge we computed the ICC(1,1) variant (Bari et al., 2019). The higher the ICC, the higher the edge contribution to subject identifiability. From now on, we will refer to this measure as “subject ICC”. Given the small sample size, a bootstrap procedure was applied when computing ICC, as suggested by Bari et al. (2019): over 100 iterations, 75% of the subjects were randomly selected, and the ICC was calculated for each edge and stored in a square symmetric matrix having edges of size N^2^, where N is the number of edges. The resulting ICC matrices over all iterations were then averaged to obtain the final subject ICC values.

We then computed ICC considering TRs as units and subjects as raters. In this “TR ICC”, the higher an edge ICC, the higher the edge contribution to distinguish between TRs regardless of subjects.

### 2.7 Statistical analysis

A permutation testing framework (Sareen et al, 2021) was employed to assess the statistical significance of the observed *Idiff* and *SR* values computed from the identifiability matrices: over 1000 iterations, subjects’ test-retest FC matrices were randomly shuffled, and the *Idiff* and *SR* were computed on the resulting randomized identifiability matrices, in order to create two non-parametric null distributions, respectively for *Idiff* and *SR*. Moreover, to correct for multiple comparisons, we merged the null distributions from all the five TRs in the within-TR analysis, and from all the TR pairwise comparisons in the between-TR analysis. The observed *Idiff* and *SR* values were then compared against their corresponding null distribution, such that the computed p-values are the proportion of values in the null distribution greater or equal to the observed values (Nichols and Holmes, 2002).

Furthermore, distributions of *Idiff*, *Iself*, *Iothers* and *SR* on a subject level (Bari et al., 2019) were calculated in order to: i) compare differential identifiability (*Idiff*) computed on the original FC matrices (before PCA) to the *Idiff* computed on the PCA-reconstructed FC matrices (both within-TR and between-TR identifiability); ii) compare FC identifiability in one TR against FC identifiability in all the other TRs (within-TR analysis); iii) compare FC identifiability in one TR comparison against FC identifiability in all the other TR pairwise comparisons (between-TR analysis); iv) compare FC identifiability computed on all volumes against FC identifiability computed only on the first 150 volumes (both within-TR and between-TR identifiability). The group distributions of *Idiff*, *Iself* and *Iothers* were compared using the Wilcoxon signed rank test followed by a False Discovery Rate (FDR) correction. Statistical differences in *SR*, instead, were assessed using the exact version of McNemar’s test for proportions in paired samples, followed by a FDR correction.

Significance threshold after FDR correction was declared at α level = 0.05. Statistical analysis was performed using R Studio (Team, R. c, 2013).

## 3. Results

### 3.1 Whole-brain within-TR connectome fingerprinting: the effect of acquisition TR and number of volumes

We first investigated how the temporal resolution at which rs-fMRI data were acquired affected the possibility to identify one individual among the whole sample only on the basis of whole-brain resting-state FC profiles (within-TR fingerprinting).

As a result of the group-level PCA, FC matrices for each subject and each TR session were reconstructed using a number of principal components (PCs) equal to the sample size (optimal number of PCs maximizing *Idiff*: m* = 20; **Table 1**; see also **Supplementary material S1**, **Supplementary figure S1A**). We found that the *SR* at which subjects were identified based on their FC was equal to 100% within all TR sessions except two (TR 0.7 s: *SR* = 97.5%; TR 2 s: *SR* = 95%). When comparing observed *SR* and *Idiff* computed on the single identifiability matrices against their correspondent null distributions (Sareen et al., 2021; see **Methods**), a statistically significant effect was obtained (permutation testing, *p* < 0.001) for each TR session. Taken together, these results suggested that, independently of TR, PCA-based reconstruction of FC ensured a successful subject identifiability (**Figure 2A**).

**Table 1.** Within-TR analysis summary table. For each TR, values of the percentage differential identifiability (*Idiff*), self-identifiability (*Iself*), others-identifiability (*Iothers*) and success rate (*SR*) are reported for the all volumes condition. Orig values were computed on the identifiability matrices derived from the original (before Principal Component Analysis reconstruction) functional connectivity matrices, while Recon values were extracted from the FC matrices reconstructed by using the optimal number of principal components (m*), for which explained variance (R^2^) is also reported.

| TR (s) | $m^*$ | $R^2$ | $Idiff_{orig}$ | $Idiff_{recon}$ | $Isself_{recon}$ | $Iothers_{recon}$ | $SR_{orig}$ | $SR_{recon}$ |
| --- | --- | --- | --- | --- | --- | --- | --- | --- |
| 0.5 | 20 | 0.84 | 38.52 | 64.19 | 96.34 | 32.15 | 100.00 | 100.00 |
| 0.7 | 20 | 0.85 | 27.32 | 46.45 | 91.15 | 44.71 | 97.50 | 97.50 |
| 1 | 20 | 0.86 | 25.57 | 44.07 | 92.83 | 48.76 | 97.50 | 100.00 |
| 2 | 20 | 0.87 | 25.20 | 43.77 | 91.44 | 47.67 | 95.00 | 95.00 |
| 3 | 20 | 0.84 | 32.03 | 56.23 | 93.36 | 37.13 | 100.00 | 100.00 |

**Figure 2.**
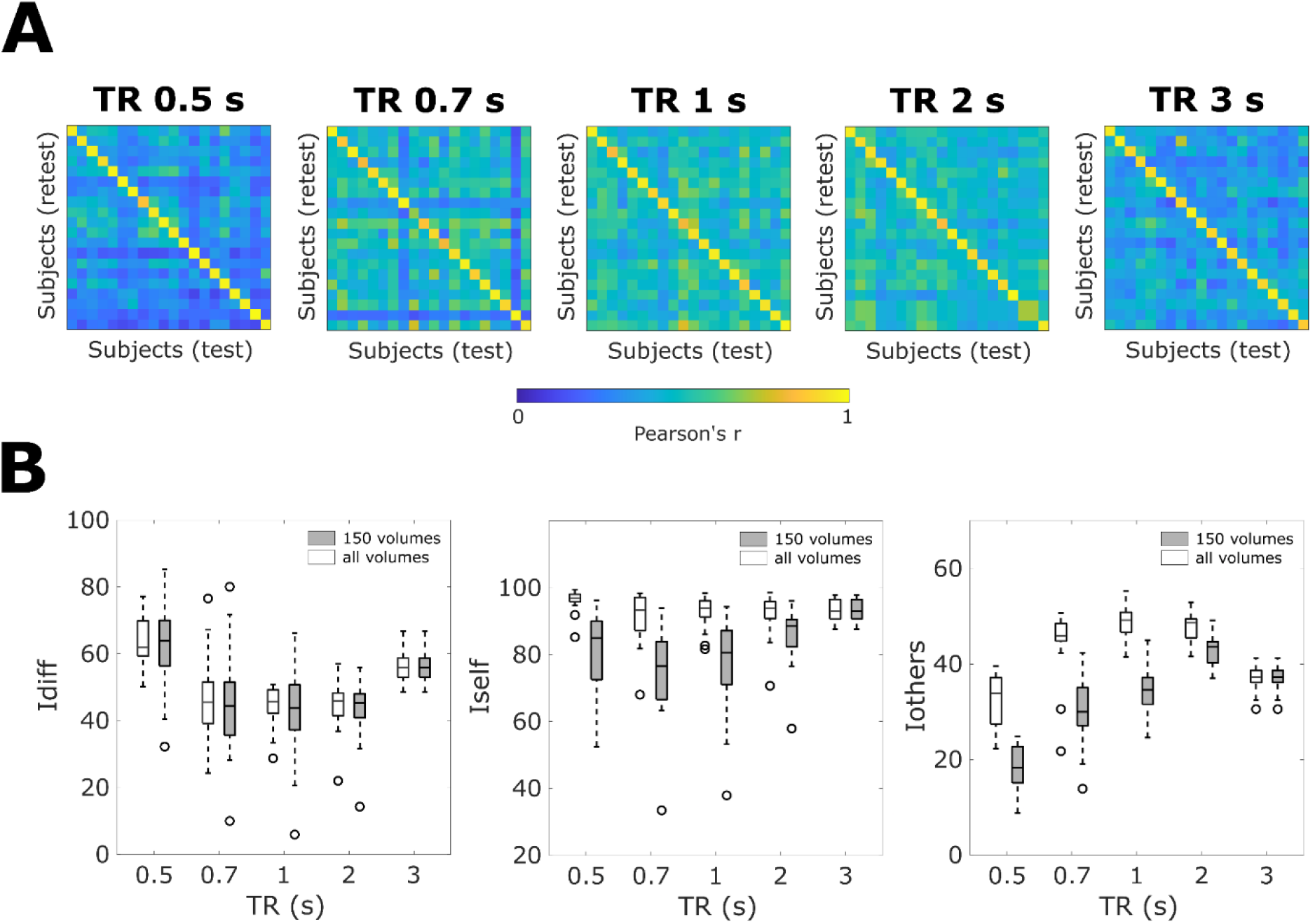
Within-TR fingerprinting. (**A**) Identifiability matrices of PCA-reconstructed functional connectivity profiles, extracted from the whole scanning time course, separately for each TR session. (**B**) Box plots of differential identifiability (*Idiff*), self-identifiability (*Iself*) and others-identifiability (*Iothers*) distributions computed at a subject-level after PCA-reconstruction, when using all (white) and 150 (gray) volumes for the fingerprinting analysis.

However, pairwise within-TR *Idiff* comparisons (**Figure 2B**) revealed that *Idiff* at TR 0.5 s was significantly higher than *Idiff* from all other TRs (*p_FDR_* < 0.05, Wilcoxon test, **Table 2**), and that TR 3 s significantly outperformed TR 0.7, 1 and 2 s in *Idiff* (*p_FDR_* < 0.05, Wilcoxon test, **Table 2**). To better explore what drove this U-shaped effect of temporal resolution in *Idiff* (**Figure 2B**), we looked at the trajectories of *Iself* and *Iothers* distributions separately. We found (**Table 2**) that at TR 0.5 s, *Iself* resulted to be significantly higher than at all other TRs. Instead, *lothers* were significantly lower at TRs 0.5 and 3 s, thus giving the inverse U-shaped trend with respect to *Idiff*. The comparisons of *SR* distributions between TRs, instead, did not reveal any significant effect.

**Table 2.** Within-TR statistics summary table. For each TR comparison, *pFDR* values are reported for the all volumes analysis (**\***: *pFDR* < 0.05, **\*\***: *pFDR* < 0.001). Moreover, for each TR, *pFDR* values resulting from the comparison between the all volumes and the 150 volumes distributions are summarized. Wilcoxon signed rank was used for differential identifiability (*Idiff*), self-identifiability (*Iself*) and others-identifiability (*Iothers*), while McNemar’s test for proportions in paired samples was employed for success rate (*SR*). Statistics refer to functional connectivity matrices reconstructed by using the optimal number of principal components.

| | | TR (s) | $p_{FDR}$ (Wilcoxon) | | | |
| --- | --- | --- | --- | --- | --- | --- |
|  |  |  | 0.5 | 0.7 | 1 | 2 |
| All volumes | ldiff | 0.7 | $1.26 \times 10^{-4} **$ | - | - | - |
| | lself | | $0.02 *$ | - | - | - |
| | lothers | | $1.91 \times 10^{-5} **$ | - | - | - |
|  | SR |  | 1.00 | - | - | - |
| | ldiff | 1 | $4.77 \times 10^{-6} **$ | 0.43 | - | - |
| | lself | | $0.04 *$ | 0.99 | - | - |
| | lothers | | $4.77 \times 10^{-6} **$ | $5.37 \times 10^{-4} **$ | - | - |
|  | SR |  | 1.00 | 1.00 | - | - |
| | ldiff | 2 | $4.77 \times 10^{-6} **$ | 0.99 | 0.43 | - |
| | lself | | $0.02 *$ | 0.99 | 0.99 | - |
| | lothers | | $4.77 \times 10^{-6} **$ | 0.17 | 0.14 | - |
|  | SR |  | 1.00 | 1.00 | 1.00 | - |
| | ldiff | 3 | $5.37 \times 10^{-4} **$ | $4.50 \times 10^{-3} *$ | $4.77 \times 10^{-6} **$ | $4.77 \times 10^{-6} **$ |
| | lself | | $0.01 *$ | 0.99 | 0.99 | 0.99 |
| | lothers | | $1.79 \times 10^{-3} *$ | $1.21 \times 10^{-3} *$ | $4.77 \times 10^{-6} **$ | $4.77 \times 10^{-6} **$ |
|  | SR |  | 1.00 | 1.00 | 1.00 | 1.00 |
| All vs 150 volumes | ldiff |  | 0.76 | 0.76 | 0.76 | 0.76 |
| | lself | | $2.29 \times 10^{-5} **$ | $1.09 \times 10^{-4} **$ | $9.54 \times 10^{-5} **$ | $7.23 \times 10^{-4} **$ |
| | lothers | | $2.54 \times 10^{-6} **$ | $2.54 \times 10^{-6} **$ | $2.54 \times 10^{-6} **$ | $9.56 \times 10^{-5} **$ |
|  | SR |  | 1.00 | 1.00 | 1.00 | 1.00 |

Similar results were found when using only the first 150 volumes instead of the whole rs-fMRI time course for the fingerprinting analysis (**Supplementary material S2, Supplementary table S1, Supplementary table S2**). No significant differences were observed in any TR condition when comparing *ldiff* and *SR* in the 150 volumes with their corresponding distributions in the analysis encompassing the whole time series (**Table 2**). *Iself* and *Iothers* values, instead, significantly decreased in all TRs (Wilcoxon test, *p_FDR_* < 0.001, **Table 2**) when a lower number of volumes was used to estimate subject identifiability.

To summarize the whole-brain within-TR rs-fMRI fingerprinting findings: i) although subject identifiability is consistently high across theTRs evaluated, it is significantly maximized at TR 0.5 and 3 s, a pattern that appears driven by lower across-subjects variability at those TRs; ii) the number of rs-fMRI volumes does not significantly affect identifiability, regardless of TR.

### 3.2 Whole-brain between-TR connectome fingerprinting: the effect of acquisition TR and number of volumes

Next, we proceeded to investigate the impact of temporal resolution on subject identifiability across different TR acquisitions. FC matrices for each TR pairwise comparison were reconstructed using a number of principal components (PCs) close to the sample size (m* = 20.7 ± 0.68, **Table 3**; see also **Supplementary material S1**, **Supplementary figure S1C**). The comparison between observed *SR* and *Idiff* against their corresponding null distributions (Sareen et al., 2021; see **Methods**) resulted in a statistically significant effect (permutation testing, *p* < 0.001). As a consequence, even if identifiability metrics showed lower values than in the within-TR analysis, PCA-based reconstruction of FC still proved to allow a successful subject identifiability.

**Table 3.**
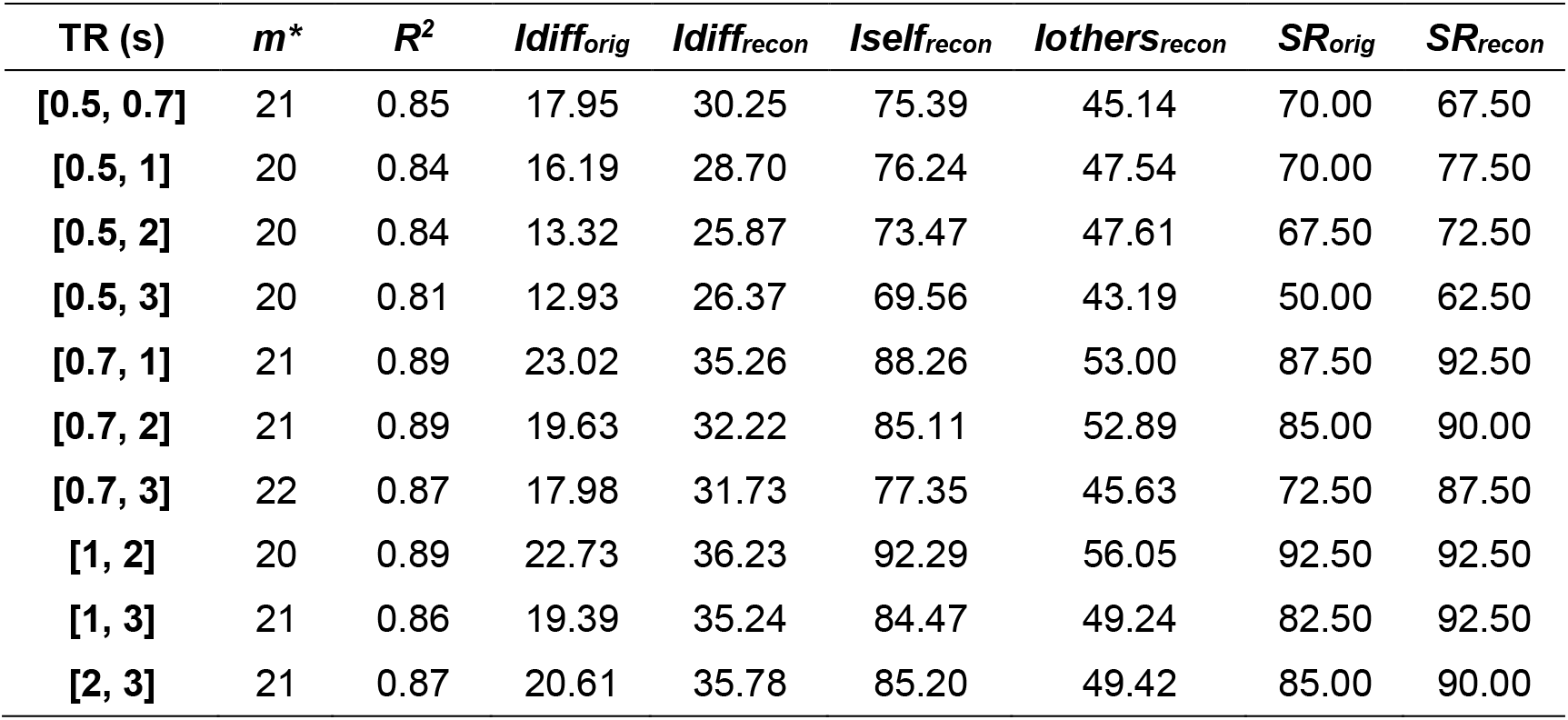
Between-TR analysis summary table. For each TR combination, values of the percentage differential identifiability (*Idiff*), self-identifiability (*Iself*), others-identifiability (*Iothers*) and success rate (*SR*) are reported for the all volumes condition. Orig values were computed on the identifiability matrices derived from the original (before Principal Component Analysis reconstruction) functional connectivity matrices, while Recon values were extracted from the FC matrices reconstructed by using the optimal number of principal components (m*), for which explained variance (R^2^) is also reported.

Additionally, we explored differences in identifiability metrics between TR pairwise combinations. Despite not always surviving FDR correction, we found a non-significant trend (*pFDR* > 0.28) according to which higher similarity between TRs resulted in higher identifiability as measured by *Idiff and SR* (**Figure 3**, **Table 4**). Moreover, *Idiff* and *SR* between close but slow TRs (i.e., 2 and 3 s) tended to be higher than between close but fast TRs (i.e., 0.5 and 0.7 s), despite the absolute difference in temporal resolution being higher in the former (1 s) rather than in the latter case (2 s) (**Figure 3B-3C**). The same trend was observed when considering the separate contributions of *Iself* and *Iothers* to subject identifiability (**Supplementary figure S4**), with p-values surviving FDR correction in several TR combinations (**Supplementary table S4**).

**Figure 3.**
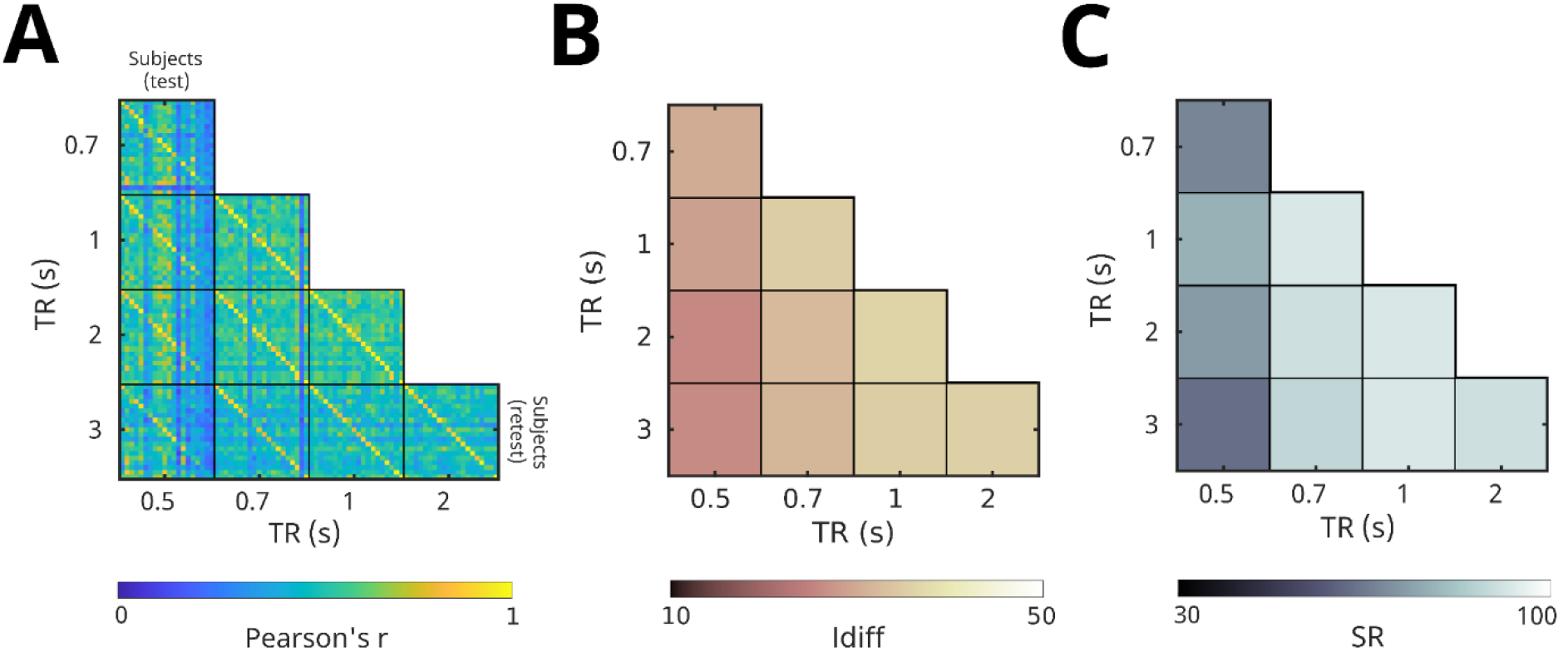
Between-TR fingerprinting. (**A**) Identifiability matrices of the PCA-reconstructed functional connectivity profiles, extracted from the whole scanning time course, separately for each TR combination. (**B**) Confusion matrices of the differential identifiability (*Idiff*) values computed from the between-TR identifiability matrices (A). (**C**) Confusion matrices of the success rate (*SR*) values computed from the between-TR identifiability matrices (A).

**Table 4.** Between-TR statistics summary table. For each comparison between TR combinations, *pFDR* values are reported for the all volumes analysis (**\***: *pFDR* < 0.05, **\*\***: *pFDR* < 0.001). Moreover, for each TR combination, *pFDR* values resulting from the comparison between the all volumes and the 150 volumes distributions are summarized. Wilcoxon signed rank was used for differential identifiability (*Idiff*), while McNemar’s test for proportions in paired samples was employed for success rate (*SR*). Statistics refer to functional connectivity matrices reconstructed by using the optimal number of principal components.

|  |  | [TR<sub>i</sub>, TR<sub>j</sub>]<br>(s) | p<sub>FDR</sub> (Wilcoxon) |  |  |  |  |  |  |  |  |  |
| --- | --- | --- | --- | --- | --- | --- | --- | --- | --- | --- | --- | --- |
|  |  |  | [0.5, 0.7] | [0.5, 1] | [0.5, 2] | [0.5, 3] | [0.7, 1] | [0.7, 2] | [0.7, 3] | [1, 2] | [1, 3] | [2, 3] |
| All<br>volumes | Idiff | [0.5, 1] | 0.57 | - | - | - | - | - | - | - | - | - |
|  | SR |  | 1.00 | - | - | - | - | - | - | - | - | - |
|  | Idiff | [0.5, 2] | 0.29 | 0.35 | - | - | - | - | - | - | - | - |
|  | SR |  | 0.75 | 1.00 | - | - | - | - | - | - | - | - |
|  | Idiff | [0.5, 3] | 0.34 | 0.78 | 0.60 | - | - | - | - | - | - | - |
|  | SR |  | 1.00 | 0.91 | 0.45 | - | - | - | - | - | - | - |
|  | Idiff | [0.7, 1] | 0.57 | 0.43 | 0.28 | 0.28 | - | - | - | - | - | - |
|  | SR |  | 0.07 | 0.16 | 0.23 | 0.07 | - | - | - | - | - | - |
|  | Idiff | [0.7, 2] | 0.71 | 0.71 | 0.34 | 0.35 | 0.28 | - | - | - | - | - |
|  | SR |  | 0.07 | 0.16 | 0.23 | 0.07 | 1.00 | - | - | - | - | - |
|  | Idiff | [0.7, 3] | 0.74 | 0.71 | 0.34 | 0.43 | 0.57 | 0.86 | - | - | - | - |
|  | SR |  | 0.07 | 0.16 | 0.23 | 0.07 | 1.00 | 1.00 | - | - | - | - |
|  | Idiff | [1, 2] | 0.43 | 0.57 | 0.28 | 0.28 | 0.90 | 0.43 | 0.60 | - | - | - |
|  | SR |  | 0.07 | 0.07 | 0.09 | 0.06 | 1.00 | 1.00 | 1.00 | - | - | - |
|  | Idiff | [1, 3] | 0.60 | 0.47 | 0.28 | 0.28 | 0.60 | 0.72 | 0.85 | 0.53 | - | - |
|  | SR |  | 0.06 | 0.07 | 0.07 | 0.06 | 0.75 | 0.75 | 0.75 | 1.00 | - | - |
|  | Idiff | [2, 3] | 0.58 | 0.53 | 0.28 | 0.28 | 0.73 | 0.28 | 0.53 | 0.70 | 0.28 | - |
|  | SR |  | 0.11 | 0.16 | 0.23 | 0.07 | 1.00 | 1.00 | 1.00 | 1.00 | 0.75 | - |
| All vs 150<br>volumes | Idiff |  | 0.12 | **0.01 \*** | **0.01 \*** | **0.01 \*** | 0.06 | **0.05 \*** | **0.01 \*** | **0.04 \*** | 0.39 | 0.39 |
|  | SR |  | 0.16 | 0.31 | 0.16 | 0.31 | 0.42 | 0.56 | 0.56 | 0.56 | 0.42 | 1.00 |

When using only the first 150 volumes, instead of the whole time course for the fingerprinting analysis, PCA-based reconstruction of FC still proved to allow a successful subject identifiability for all pairwise TR combinations (see **Supplementary material S3**). However, relative to the use of the full time series, using only the first 150 rsfMRI volumes had some identifiability costs: the variability of *ldiff* was higher (**Table 3**), and *SR* was on average lower (68.5% ± 20.35) with respect to the full time time course (82.5% ± 11.49). Notably, when comparing the *Idiff* and *SR* of each TR pairwise combination in the 150 volumes with their corresponding values in the all volumes analysis, we found that, when using less timepoints, *Idiff* was significantly lower in most TR combinations involving fast TRs (i.e., TR 0.5 and TR 0.7 s), while no *SR* value comparison survive FDR correction (**Table 4**). A significant decrease of *lself* and *lothers* was observed in the 150 volumes condition (**Supplementary table S6**) for most TR combinations.

To summarize the whole-brain across-TR fingerprinting findings: i) subject identifiability tends to be higher for rs-fMRI acquisition protocols using similar TRs; ii) using the first 150 rs-fMRI volumes decreases identifiability performance with respect to the use of the full acquired time series for each TR.

### 3.3 Local contributions to rs-fMRI fingerprinting: identifiability of subjects across TRs and identifiability of TRs across subjects

The within- and between-TR analysis discussed in the previous paragraphs referred to the whole brain FC profiles. To explore whether specific edges or brain regions contributed more than others to subject identifiability over all TRs, we computed a subject ICC. We found that within- and between-network connections involving high-level associative areas – that is, brain regions belonging to fronto-parietal, Default Mode Network, dorsal and ventral attention networks (Damoiseaux et al., 2006) – had a prominent role in separating between subjects, regardless of TR (**Figure 4A-4C-4E**). As an opposite pattern with respect to subject ICC, low-level areas – comprising somatomotor and subcortical regions (Damoiseaux et al., 2006; Xu et al., 2020) – appear among the top 5 networks highly implicated in distinguishing between TRs, regardless of subjects (**Figure 4B-4D-4F**).

Finally, it is interesting to note that both subject and TR ICC results did not qualitatively change much when rerunning the analysis taking into account only the first 150 volumes of each scanning session (**Supplementary figure S6**).

**Figure 4.**
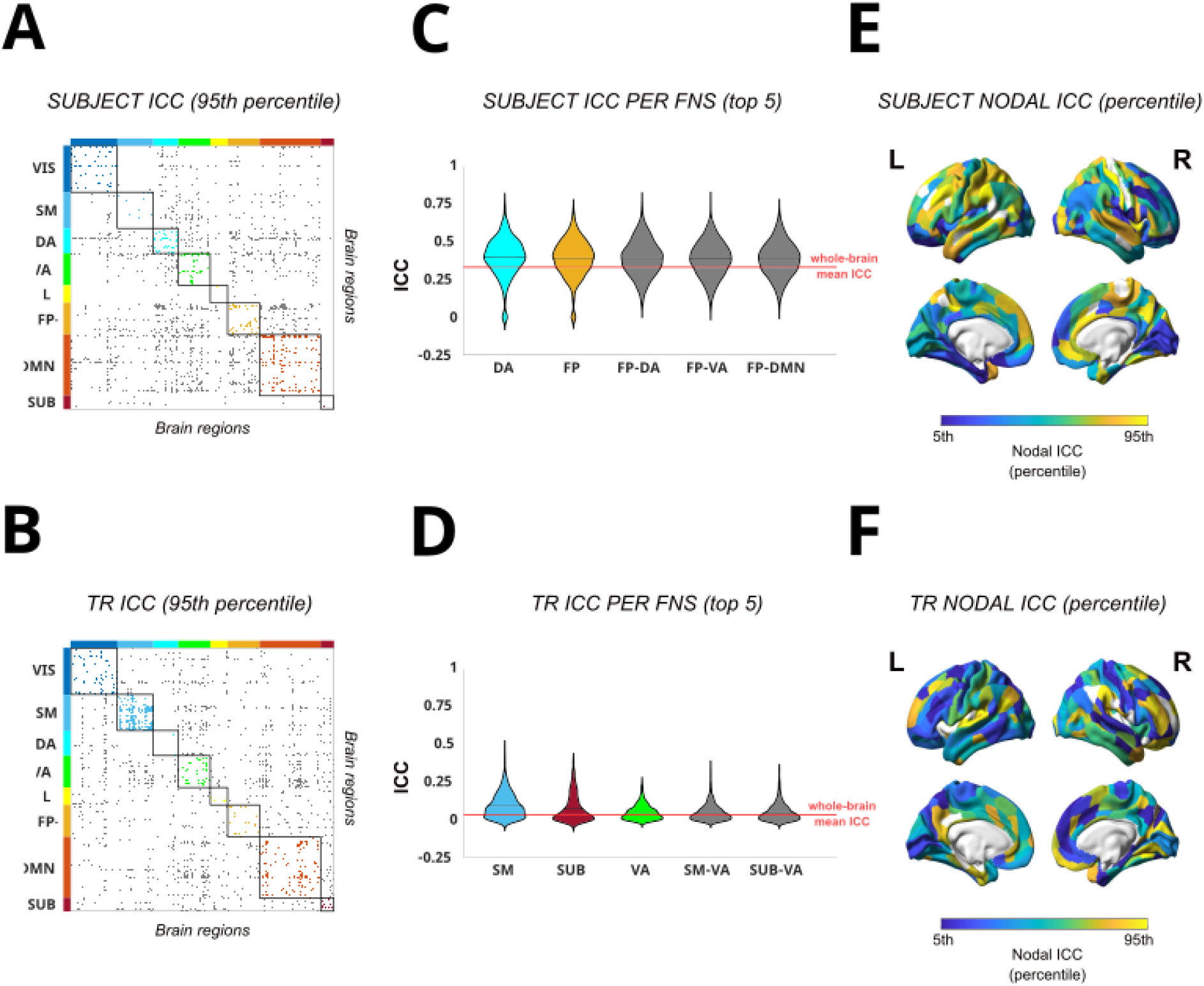
Edgewise Intra-class correlation (ICC) analysis of subject identifiability and task identifiability. (**A-B**) Edgewise subject (A) and TR ICC (B) matrices, showing only functional connections with ICC values significantly higher than the mean distribution (that is, lying in the 95th percentile). The brain regions are ordered according to Yeo’s (Yeo et al., 2011) functional resting state networks (FNs): Visual (VIS), Somato-Motor (SM), Dorsal Attention (DA), Ventral Attention (VA), Limbic system (L), Fronto-Parietal (FP), Default Mode Network (DMN), and subcortical regions (SUB). The colored dots refer to within FNs networks edges, while gray dots refer to between FNs networks edges, as in Amico & Goñi (2018). (**C-D**) Violin plots of edgewise subject (C) and TR (D) ICC distributions for the 5 FNs with the highest mean ICC value. Each colored violin plot indicates a different within FN, while gray violin plots indicate between FNs ICC distributions. The horizontal solid black line within each violin plot indicates the mean value of each distribution; the solid red line across the violin plots, instead, indicates the whole-brain mean ICC value, as in Amico & Goñi (2018). The most prominent FNs for subject’s identifiability (C) resulted: DA, FP, and the DA-FP, FP-DMN, FP-VA interactions. For TR identifiability (D) the most relevant FNs were: SM, SUB, VA and the VA-SUB, SM-VA interactions. (**E-F**) Brain render of nodal ICC, computed as the column-wise mean of the edgewise ICC matrices for both subject (E) and TR (F) ICC, and represented at 5th – 95th percentile threshold. Nodal ICC gives an assessment of the overall prominence of each brain region for subject’s and TR identifiability. All plots refer to the all volumes analysis.

## 4. Discussion

In this study, we employed brain connectome fingerprinting enriched by group-level principal component analysis to assess subject identifiability across resting-state fMRI protocols with varying temporal resolution. This allowed us to dig into the still unresolved question of test-retest reliability in fast fMRI approaches (Jahanian et al., 2019; Polimeni & Lewis, 2021), and to make specific recommendations about pooling data acquired with different BOLD signal sampling rates, which has useful applications in neuroscientific research, in both clinical and computational neuroimaging. In particular, we provide compelling evidence that (i) when comparing acquisitions with the same TR, subject identifiability was successful no matter the TR tested (0.5, 0.7, 1, 2, 3 s), but it was maximized at TR 0.5 and 3 s; (ii) when comparing acquisitions with different TRs, the more similar the TRs, the higher the subject identifiability, which was especially true for slow TRs (i.e., TR 2 and 3 s); (iii) the number of acquisition timepoints (either full time course or the first 150 volumes) affected the between-TR but not the within-TR analysis; (iv) higher-level cognitive brain areas contributed significantly more to subject identifiability, while lower-level cognitive regions played a crucial role in distinguishing between TRs.

We replicated the finding that intrinsic functional connectivity is unique and sufficient to identify an individual among a sample of healthy subjects (Amico & Goñi, 2018; Bari et al., 2019; Finn et al., 2015; Romano et al., 2022; Sareen et al., 2021; Sorrentino et al., 2021; Svaldi et al., 2021; Troisi Lopez et al., 2022; Van de Ville et al., 2021). We further extended these results by proving that, despite the high identifiability achieved regardless of the temporal resolution at which data were acquired, fingerprinting analysis was still sensitive to the fMRI acquisition protocol, and in particular to TR.

### 4.1 Within-TR subject identifiability is maximized at TR 0.5 and 3 s

Although in all within-TR sessions subject identifiability was successfull, we showed that Idiff across scanning sessions with the same TR was maximized at TR 0.5 s and 3 s, while significantly decreasing at TR 0.7, 1 and 2 s. These findings can be explained by considering that test-retest reliability of resting state fMRI data is largely affected by the noise introduced by scanner artifacts, subject movements, changes in cognitive or emotional states, and, importantly, by physiological factors (Caballero-Gaudes & Reynolds, 2017). Cardiorespiratory brain pulsations are temporally correlated with the BOLD fMRI signal, which means that in typical rs-fMRI protocols with TRs ∼2 s, these confounding components are added up to and can mask the intrinsic neural signal (Özbay et al., 2019). In fact, FC BOLD fluctuations predominantly lie below 0.1 Hz frequencies (Cordes et al., 2001), thus a low-pass filter is commonly applied to retain only frequencies below this threshold, and to remove signal from non-neuronal sources, such as respiratory rate (0.2-0.3 Hz; Kiviniemi et al., 2016) and cardiac pulsations (0.8-1.2 Hz; Kiviniemi et al., 2016). However, this denoising procedure is efficient only when TR is short (< 0.5 s; Jahanian et al., 2019; Liu, 2016; Reynaud et al., 2017), reducing the noise components in the signal and thus improving test-retest reliability (Jahanian et al., 2019). Instead, when TR is higher than 0.5 s, cardiac fluctuations are undersampled and aliased into the lower frequency range, which makes physiological noise difficult to remove with band-pass filtration (Lowe et al., 1998; Houtari et al., 2019). The consequent increase of aliased noise in the signal is consistent with the significantly lower differential identifiability we registered in TR 0.7, 1 and 2 s sessions. Finally, at TR 3 s, according to the Nyquist theorem, the BOLD signal is sampled at ∼0.16 Hz, implicitly low-pass filtering respiratory and cardiovascular noise (Houtari et al., 2019), which is consistent with the improved test-retest reliability. A simulation is included in the **Appendix** to further support this explanation.

### 4.2 Between-TR subject identifiability is maximized when TRs are more similar

Subject identifiability was successful even when considering rs-fMRI acquisitions with different TRs. However, identifiability gradually decreased as TRs became more dissimilar. This confirms the intuition that the more similar the signal sampling, the higher the test-retest reliability. We expanded on this quite trivial finding by showing that identifiability between close but slow TRs (i.e., 2 s and 3 s) tended to be higher than identifiability between close but fast TRs (i.e., 0.5 s and 0.7 s), despite the absolute difference in temporal resolution being higher in the former (1 s) rather than in the latter case (0.2 s). In fact, previous results underlined that physiological (cardiac and respiratory) and low frequency fluctuations reflecting functional connectivity exhibited maps increasingly correlated as a function of TR increase, both when considering spatial power and frequency power analysis (Houtari et al., 2019). As a consequence, at close but slow TRs noise is aliased into neural frequency more similarly than in close but fast TRs, which leads to a higher test-retest reliability, even though not significantly different.

### 4.3 The number of rs-fMRI volumes affects between-TR but not within-TR subject identifiability

Our experimental design used a constant acquisition time for each rs-fMRI session (7.4 min), which led to different number of brain volumes (NoV) for each temporal resolution (TR(s)/NoV = 0.5/905; 0.7/646; 1/452; 2/226; 3/150). We investigated the effect of number of volumes on identifiability by using the full time series of each protocol and the first 150 volumes. In both within- and between-TR analysis, subjects could be correctly identified even when a limited number of volumes was used, in agreement with previous literature (Amico & Goñi, 2018). However, when comparing sessions with different TRs, *Idiff* computed on the whole time course was significantly higher than when considering only a subset of volumes. This supports prior evidence that reproducibility improves with longer scanning length, and thus with a larger amount of data (Anderson et al., 2011; Birn et al., 2013; Noble et al., 2017, Van de Ville et al., 2021). When comparing same temporal resolution sessions, instead, the number of volumes did not have any significant impact on *Idiff*. In other words, connectome fingerprinting enriched by group-level PCA was robust enough to guarantee a good test-rest reliability regardless of TR and number of volumes here tested, and, in the most extreme case, even across the equivalent of two acquisitions lasting 1.25 s each at TR 0.5 s (150 volumes). Our findings contradict previous studies suggesting that long scanning acquisitions, ranging from 36 (Noble et al., 2017) up to 90 minutes (Laumann et al., 2015), are needed to reach functional connectivity reliability. In addition, our results also challenge the evidence that about 5-10 min of data are sufficient to ensure a stable measurement of connectivity (Choe et al., 2015; Jahanian et al., 2019; Shehzad et al., 2009; Tomasi et al., 2017; Van Dijk et al., 2010). Instead, our study shows that, even with very short scanning duration, the richness of information coming with higher temporal resolution, and thus higher number of acquired timepoints, guaranteed a test-retest reliability as good as when data were acquired with standard protocols (∼2-3 TRs). This is consistent with previous findings generated by randomly (Shah et al., 2016) or systematically (Birn et al., 2013; Houtari et al., 2019) downsampling BOLD signal, and by employing feedforward neural networks on an even more limited number of timepoints in fast fMRI acquisitions (TR(s)/NoV = 0.72/100, Sarar et al., 2020).

### 4.4 The contribution of specific brain networks to edgewise connectivity to rs-fMRI fingerprinting

In addition to assessing fingerprinting at whole-brain level, we employed ICC to investigate whether some specific edges will play a differential role in making each individual’s functional profile unique. We found that connections within and between fronto-parietal, Default Mode Network, dorsal and ventral attention networks (classically classified as associative areas, Damoiseaux et al., 2006) were the main drivers of subject identifiability. On the contrary, edges belonging to visual, somatomotor and limbic networks gave poor contribution to fingerprinting. This was in line with prior results (Amico & Goñi, 2018; Finn et al., 2015; Van de Ville et al., 2021), but we extended this result by showing that this effect was not impacted by the variability in the temporal resolution of the rs-fMRI acquisition protocols.

In order to explain this resting state networks’ specific role in subject identifiability, a link can be speculated with the intrinsic neural activity profile and the cognitive processes characterizing these brain circuits (Van de Ville et al., 2021). Previous studies concurred to distinguish between two sets of cortical regions. On one hand, visual, auditory, somatosensory and motor cortices are mainly responsible for detecting real-time rapid and potentially dangerous changes in the environment. On the other hand, frontal, temporal and parietal regions, are involved in sustain activity related to working memory (Zylberberg & Strowbridge, 2017), decision-making (Gold & Shadlen, 2007), hierarchical reasoning (Sarafyazd & Jazayeri, 2019), language (Binder et al., 2009), emotion regulation (Laird et al., 2011), and, in general, cognitive processes which require a multimodal integration of information. This differentiation is supported by evidence about synaptic receptor and ion channel gene expression (Cioli et al., 2014), cytoarchitectonic profiles (Gao et al., 2020), gray matter myelination gradients (Huntenburg et al., 2017), functional and anatomical network properties (see Mesmoudi et al., 2013), behavioral gradients (Margulies et al., 2016), and notably, neuronal time scales, with somatomotor regions exhibiting short firing patterns, and associative areas showing longer intrinsic time scales (Gao et al., 2020; Hacker et al., 2017; Runyan et al., 2017).

A relationship between these gradients and human brain identifiability can be hypothesized, by considering that low-level sensory-motor areas (as well as limbic regions) are involved in the fast processing of potentially dangerous information coming from the external environment. Thus, they code for a primitive and almost merely evolutionary function, common to all human (and non-human) brains. As such, no fundamental role in distinguishing between single individuals can be attributed to these circuits (Van de Ville et al., 2021).

On the contrary, subject connectome uniqueness mainly resides in high-level associative areas, where functional patterns are more decoupled, and thus less predictable on the basis of the structural organization of brain anatomy (Preti & Van de Ville, 2019). Moreover, interesting insights might come in the future by investigating the relationship between myelin gradients (higher myelin content in somatomotor that in associative areas, Huntenburg et al., 2017) and brain plasticity (Bonetto et al., 2020): frontal areas are among the last regions to the brain where myelin-associated factors contribute to closing the critical period in the transition between adolescence and adulthood (Crews et al., 2007), and play a fundamental role in functional connectivity fingerprinting during early brain development (Hu et al., 2022). It is intriguing to think that areas which make us unique are the ones more likely to be affected by experience-based learning and more responsible for reasoning, goal setting and emotional regulation, but these are only speculations which need fact-based and data-driven investigation (St-Onge et al., 2023).

ICC also allowed us to explore the role of each edge to distinguish between different temporal resolution sessions. An opposite pattern with respect to edgewise individual identifiability emerged: connections within and between sensory motor areas were the main drivers, while associative regions poorly contributed to separate between TRs. These findings can be explained by considering that low-level cognition areas are characterized by a brief transient neural activity which is registered by faster but not by slower TRs. Intrinsic time scales in high-level circuits, instead, are longer, can be detected in a robust way from both fast and slow TRs, and thus no temporal resolution-specific profiles can be extracted at an edgewise level in these areas.

In conclusion, we replicated previous fingerprinting results (Amico & Goñi, 2018; Finn et al., 2015; Van de Ville et al., 2021) distinguishing two sets of brain networks which provide different contributions to identifiability. These findings are also consistent with studies investigating the temporal organization of large-scale brain activity, and assigning a key role to dynamic spontaneous transitions between two metastates, a sensorimotor/perceptual and a cognitive/associative one (Vidaurre et al., 2017). Our work extends this notion by emphasizing how these metastates affect subject and TR identifiability, and how this identifiability is affected by the fMRI temporal resolution.

### 4.5 Limitations and future directions

This study has a number of limitations. First, our analysis employed the cortical Glasser multimodal parcellation atlas (Glasser et al., 2016) together with subcortical gray matter parcels from the HCP atlas. Since the choice and resolution of parcellation atlas can affect between-subject and within-session differences in FC estimations (Ahrends et al., 2022; Pervaiz et al., 2020), future studies with different parcellations are needed to further validate our findings. Second, we did not collect peripheral physiological measures such as cardiac and respiration activity, to verify a uniform physiological state of the subjects across acquisitions with different TRs. Moreover, we found that brain associative areas gave the highest contribution to fingerprinting regardless of TR. However, Van de Ville et al. (2021) showed that “burst of identifiability” can be detected at fast timescales also in low-level regions. Future work should explore the relationship between fMRI acquisition temporal resolution and fingerprinting by employing a sliding windows approach (Van de Ville et al., 2021) or other models assessing dynamic functional connectivity (Keilholz et al., 2017; Preti et al., 2017). Finally, our study only evaluates resting-state fMRI data. Applying the same framework to task-based sessions could provide interesting insights on whether functional connectomes derived by different TR sessions can explain different cognitive dimensions (Sareen et al., 2021). In fact, it has been proven that the temporal scales of fingerprinting can be linked to behavior, with faster time scales more related to, for instance, multisensory stimulation and visuospatial attention, and slower time scales more related, for example, to language, social cognition and working memory (Van de Ville et al., 2021). Following this rationale, subject identifiability might be more or less successfully achieved according to the different tasks subjects perform in the MRI scanner (Amico & Goñi, 2018; Finn et al., 2015; Van de Ville et al., 2021), and to the protocols’ temporal resolution.

## 5. Conclusions

This study demonstrates the robustness and reliability of FC fingerprinting derived from rs-fMRI across a wide range of temporal resolutions, including “fast” (TR 0.5 s) and “slow” (TR 3 s) full brain coverage, with intermediate values within the same group of healthy volunteers. Interestingly, while always high, subject identifiability was influenced by the temporal resolution. In fact, our work shows that the fastest and slowest TRs improved identifiability relative to the other intermediate TRs, which is consistent with a relatively lower contribution of physiological noise. The study also confirms that FC fingerprints have different timescales across the brain. In particular, high-level associative areas being more stable across protocols with different TRs, contributed more to subject identifiability. Instead, low-level sensorimotor networks were mostly responsible for differentiating subjects between different TR acquisitions. Overall, these findings suggest that pooling rs-fMRI data for large-sample fingerprinting analyses is feasible, but TR differences should be taken into consideration to reduce identifiability biases and to understand the contribution of different functional networks to the fingerprints and related biomarkers.

## Acknowledgements

This study was supported by the Autonomous Province of Trento, Italy (Project: “NeuSurPlan and integrated approach to neurosurgery planning based on multimodal data”), the ISMRM Exchange Award 2021-2022 “Investigating in-vivo human brain dynamic connectivity with fast fMRI”, and the Dipartimento di Eccellenza project 2018-2022 (Italian Ministry of Education, University and Research). EA acknowledges financial support from the SNSF Ambizione project “Fingerprinting the brain: network science to extract features of cognition, behavior and dysfunction” (grant number PZ00P2_185716). All the authors declare they have no competing interests.

