## Supplementary materials and Appendix for "TR(acking) individuals down: exploring the effect of temporal resolution in resting-state functional MRI fingerprinting"

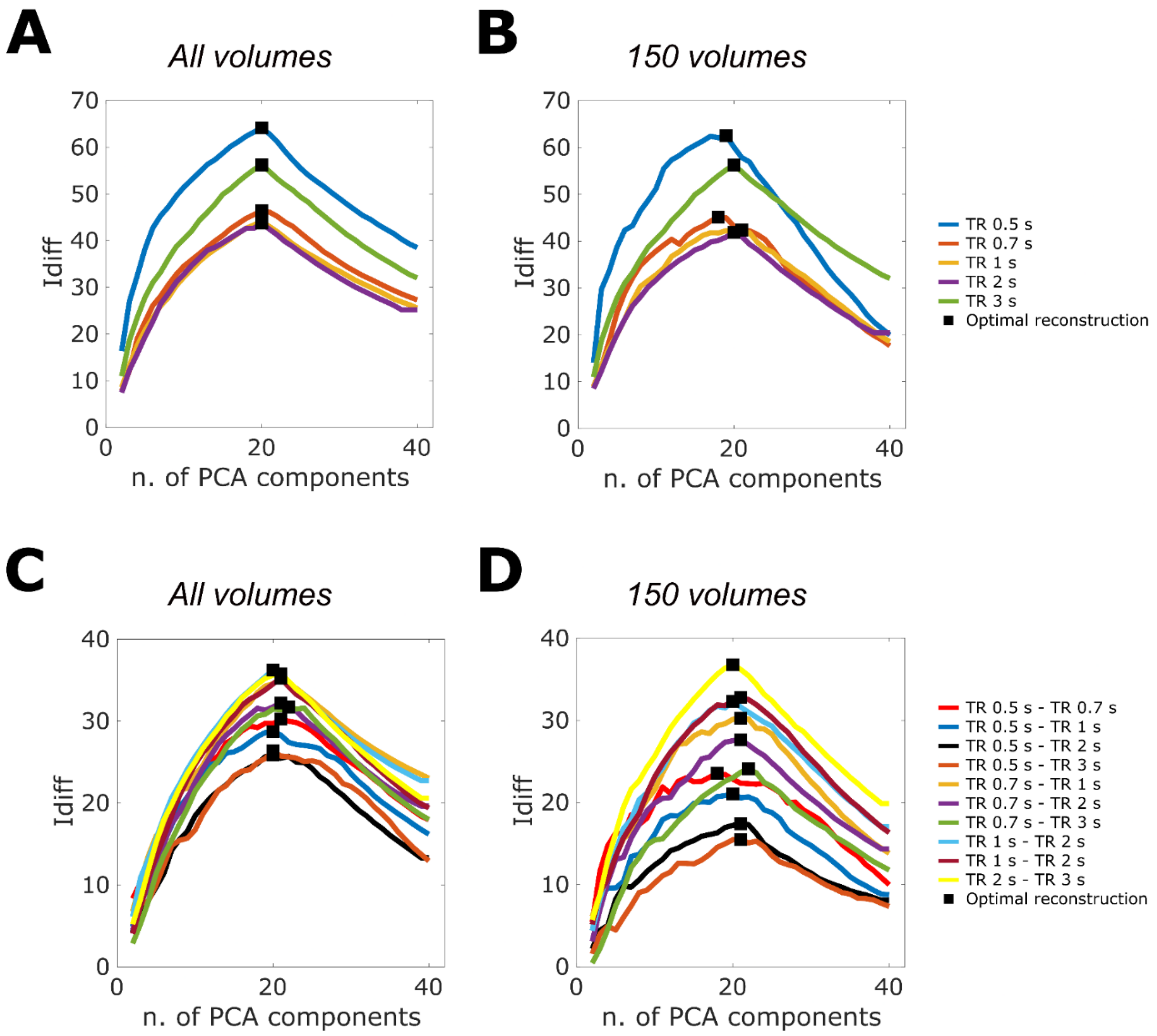


**Supplementary figure S1. Optimal number of principal components (PCs).** *Idiff* as a function of the number of PCs used to reconstruct the functional connectivity matrices in the within-TR (A-B) and between-TR (C-D) analysis, when using all volumes (A-C) or only the first 150 volumes (B-D) of the resting state fMRI session time course.

### S1. Whole-brain connectome fingerprinting: the effect of PCA reconstruction

Regardless of the fingerprinting analysis (within- or between-TR) and the number of volumes (whole time course or only the first 150 volumes), we found that *Idiff* was significantly higher after the optimal reconstruction than before PCA ($p_{FDR} < 0.001$, Wilcoxon test, **Supplementary figure S2, Supplementary figure S3**). Thus, our statistical analysis confirmed the efficacy of the data-driven denoising procedure proposed by Amico & Goñi (2018) in significantly improving identifiability at the whole brain level. Indeed, we found that by effectively eliminating group-level principal components that did not contribute to test-retest reliability, we achieved a substantial improvement of subject identifiability across sessions relative to the original ones (Amico & Goñi, 2018; Bari et al., 2019; Svaldi et al., 2021).

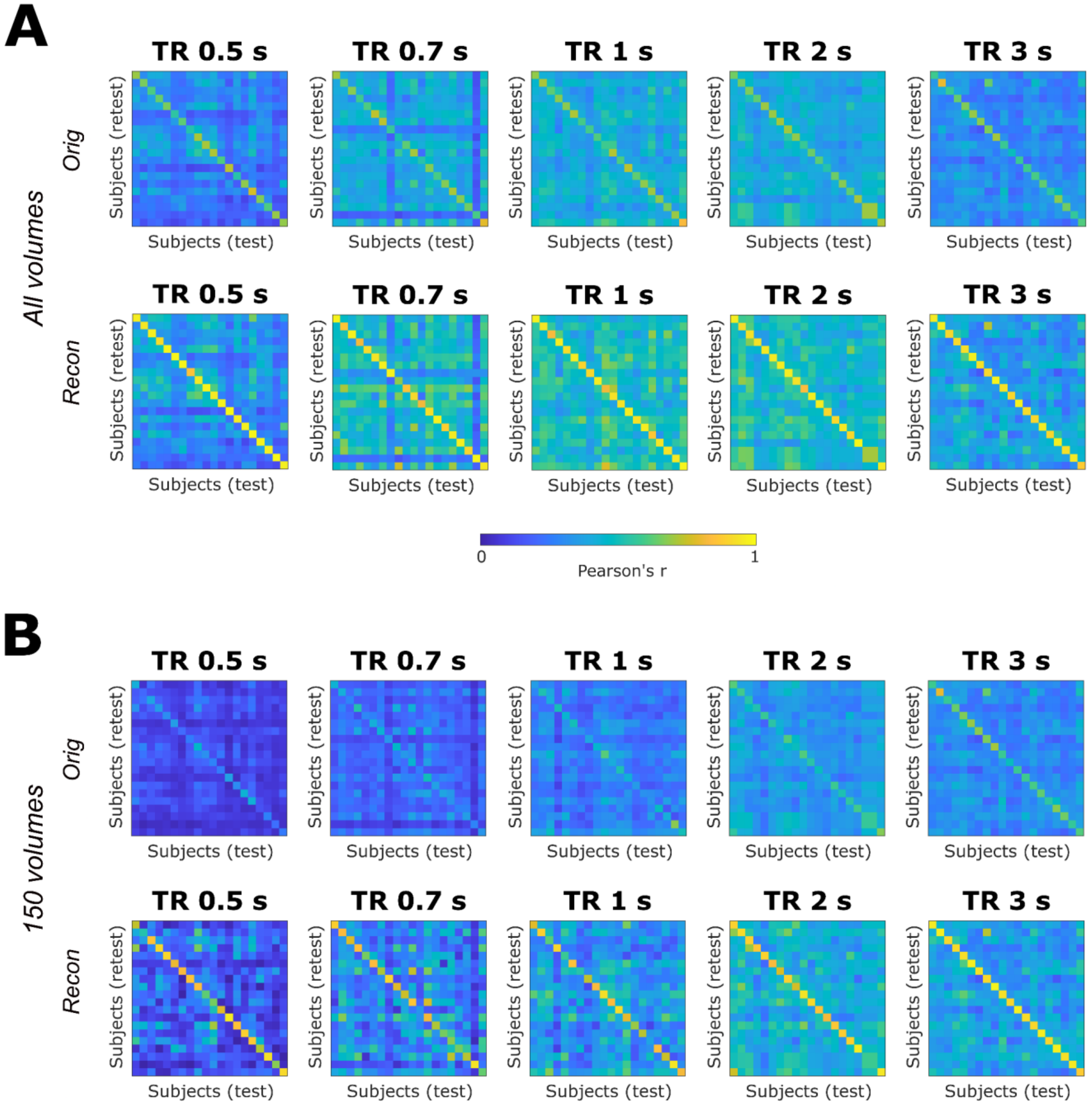


**Supplementary figure S2. Effect of PCA on within-TR fingerprinting.** Identifiability matrices of the original (*Orig*) and PCA-reconstructed (*Recon*) functional connectivity profiles, extracted from the whole time course (A) and from the first 150 volumes of the time course only (B), separately for each TR session.

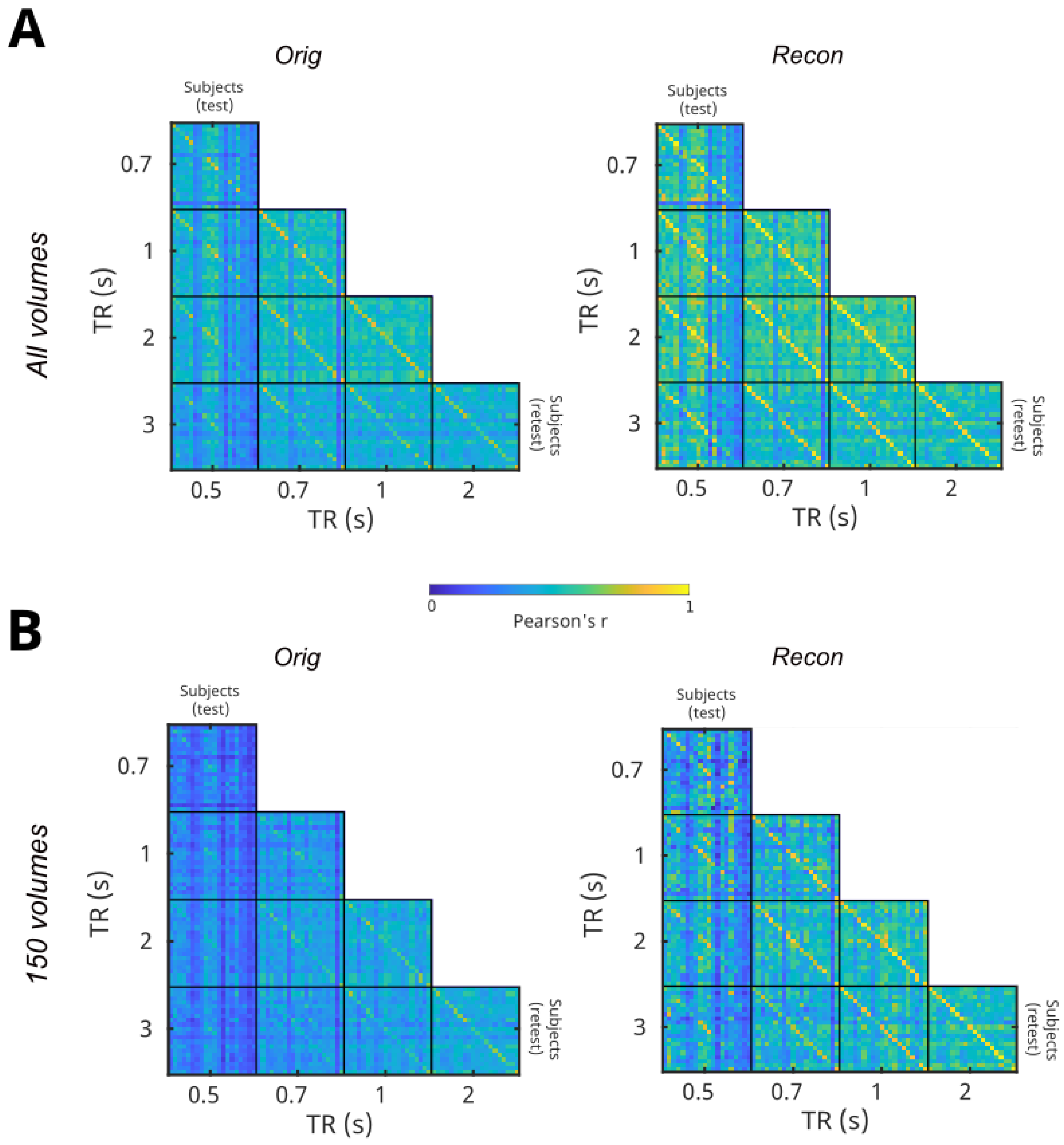


**Supplementary figure S3. Effect of PCA on between-TR fingerprinting.** Identifiability matrices of the original (*Orig*) and PCA-reconstructed (*Recon*) functional connectivity profiles, extracted from the whole time course (A) and from the first 150 volumes of the time course only (B), separately for each TR comparison.

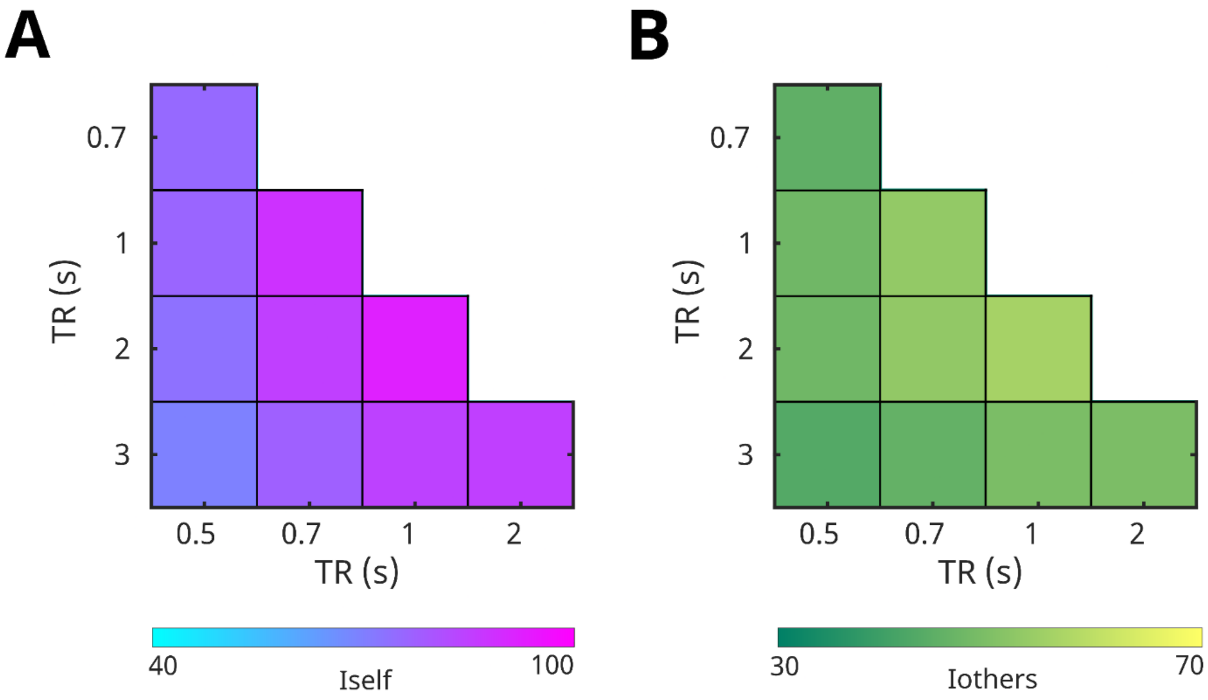


**Supplementary figure S4. All volumes between-TR fingerprinting - *Iself* and *Iothers*.** (**A**) Confusion matrices of the *Iself* values computed from the between-TR identifiability matrices (**Figure 3A**). (**B**) Confusion matrices of the *Iothers* values computed from the between-TR identifiability matrices (**Figure 3A**).

### S2. Whole-brain within-TR connectome fingerprinting: results on 150 volumes

PCA reconstruction of the FC matrices revealed an optimal number of PCs close to the sample size (m* = 19.6 ± 1.1; **Supplementary table S1**, **Supplementary figure S1B**), despite being more variable with respect to the all volumes condition. *SR* remained high (93% ± 5.4), and when comparing observed *SR* and *Idiff* computed on the single identifiability matrices against their correspondent null distributions (Sareen et al., 2021), a statistically significant effect was obtained (permutation testing, p < 0.001) for each TR session.

However, pairwise within-TR *Idiff* comparisons revealed that mean *Idiff* and *Iself* in TR 0.5 s and TR 3 s conditions were significantly higher than in all other TR sessions (Wilcoxon test, pFDR < 0.05; **Supplementary table S2**), while *SR* did not show differences between TRs, and *Iothers* distributions revealed a significant inverse U-shaped trend as described for the all volumes analysis (**Figure 2B**).

| | TR (s) | *m** | $R^2$ | $Idiff_{orig}$ | $Idiff_{recon}$ | $Iself_{recon}$ | $Iothers_{recon}$ | $SR_{orig}$ | $SR_{recon}$ |
|---|---|---|---|---|---|---|---|---|---|
| **150 volumes** | **0.5** | 19 | 0.67 | 19.95 | 62.54 | 80.77 | 18.23 | 92.50 | 95.00 |
| | **0.7** | 18 | 0.71 | 17.65 | 45.13 | 75.02 | 29.89 | 77.50 | 87.50 |
| | **1** | 21 | 0.78 | 18.54 | 42.34 | 76.81 | 34.47 | 80.00 | 87.50 |
| | **2** | 20 | 0.82 | 20.40 | 41.85 | 84.74 | 42.89 | 90.00 | 95.00 |
| | **3** | 20 | 0.84 | 32.03 | 56.23 | 93.36 | 37.13 | 100.00 | 100.00 |

**Supplementary table S1. 150 volumes within-TR analysis summary table.** For each TR, values of the percentage differential identifiability (*Idiff*), self-identifiability (*Iself*), others-identifiability (*Iothers*) and success rate (*SR*) are reported for the 150 volumes condition. *Orig* values were computed on the identifiability matrices derived from the original (before Principal Component Analysis reconstruction) functional connectivity (FC) matrices, while *Recon* values were extracted from the FC matrices reconstructed by using the optimal number of principal components (m*), for which explained variance ($R^2$) is also reported.

| | | $p_{FDR}$ (Wilcoxon) | | | |
|---|---|---|---|---|---|
| | **TR (s)** | **0.5** | **0.7** | **1** | **2** |
| Idiff | **0.7** | **5.43 x $10^{-3}$ *** | - | - | - |
| Iself | | 0.33 | - | - | - |
| Iothers | | **4.77 x $10^{-6}$ **** | - | - | - |
| SR | | 1.00 | - | - | - |
| Idiff | **1** | **2.54 x $10^{-3}$ *** | 0.33 | - | - |
| Iself | | 0.55 | 0.81 | - | - |
| Iothers | | **4.77 x $10^{-6}$ **** | **4.69 x $10^{-3}$ *** | - | - |
| SR | | 1.00 | 1.00 | - | - |
| Idiff | **2** | **4.10 x $10^{-4}$ **** | 0.63 | 0.93 | - |
| Iself | | 0.49 | **0.03 *** | 0.11 | - |
| Iothers | | **4.77 x $10^{-6}$ **** | **7.63 x $10^{-6}$ **** | **3.82 x $10^{-5}$ **** | - |
| SR | | 1.00 | 1.00 | 1.00 | - |
| Idiff | **3** | 0.1 | **0.01 *** | **5.60 x $10^{-4}$ **** | **3.82 x $10^{-5}$ **** |
| Iself | | **8.06 x $10^{-4}$ **** | **1.91 x $10^{-5}$ **** | **1.91 x $10^{-5}$ **** | **8.06 x $10^{-4}$ **** |
| Iothers | | **4.77 x $10^{-6}$ **** | **7.87 x $10^{-5}$ **** | 0.06 | **2.23 x $10^{-5}$ **** |
| SR | | 1.00 | 1.00 | 1.00 | 1.00 |

**Supplementary table S2. 150 volumes within-TR statistics summary table.** For each TR comparison, *pFDR* values are reported for the 150 volumes analysis (*****: *pFDR* < 0.05, ******: *pFDR* < 0.001). Wilcoxon signed rank was used for differential identifiability (*Idiff*), self-identifiability (*Iself*) and others-identifiability (*Iothers*), while McNemar's test for proportions in paired samples was employed for success rate (*SR*). Statistics refer to functional connectivity matrices reconstructed by using the optimal number of principal components.

| | | [TR*i*, TR*j*] (s) | [0.5, 0.7] | [0.5, 1] | [0.5, 2] | [0.5, 3] | [0.7, 1] | [0.7, 2] | [0.7, 3] | [1, 2] | [1, 3] | [2, 3] |
|---|---|---|---|---|---|---|---|---|---|---|---|---|
| | | | $p_{FDR}$ (Wilcoxon) | | | | | | | | | |
| All volumes | Iself | [0.5, 1] | 0.83 | - | - | - | - | - | - | - | - | - |
| | Iothers | | 3.44 x $10^{-3}$ * | - | - | - | - | - | - | - | - | - |
| | Iself | [0.5, 2] | 0.24 | 0.20 | - | - | - | - | - | - | - | - |
| | Iothers | | 0.01 * | 0.90 | - | - | - | - | - | - | - | - |
| | Iself | [0.5, 3] | 0.11 | 0.16 | 0.18 | - | - | - | - | - | - | - |
| | Iothers | | 0.04 * | 1.32 x $10^{-5}$ ** | 0.21 | - | - | - | - | - | - | - |
| | Iself | [0.7, 1] | 0.03 * | 0.08 | 0.03 * | 0.02 * | - | - | - | - | - | - |
| | Iothers | | 1.32 x $10^{-5}$ ** | 2.26 x $10^{-5}$ ** | 1.61 x $10^{-5}$ ** | 0.30 | - | - | - | - | - | - |
| | Iself | [0.7, 2] | 0.20 | 0.29 | 0.08 | 0.03 * | 0.99 | - | - | - | - | - |
| | Iothers | | 1.32 x $10^{-5}$ ** | 7.40 x $10^{-4}$ ** | 1.53 x $10^{-3}$ * | 2.26 x $10^{-5}$ ** | 8.58 x $10^{-6}$ ** | - | - | - | - | - |
| | Iself | [0.7, 3] | 0.71 | 0.99 | 0.55 | 0.18 | 1.50 x $10^{-3}$ * | 0.01 * | - | - | - | - |
| | Iothers | | 0.90 | 6.30 x $10^{-4}$ ** | 5.35 x $10^{-4}$ ** | 1.61 x $10^{-5}$ ** | 0.67 | 0.02 * | - | - | - | - |
| | Iself | [1, 2] | 0.02 * | 0.01 * | 1.50 x $10^{-3}$ * | 1.41 x $10^{-3}$ * | 0.11 | 0.01 * | 4.29 x $10^{-4}$ ** | - | - | - |
| | Iothers | | 8.58 x $10^{-6}$ ** | 0.18 | 0.18 | 0.15 | 1.32 x $10^{-5}$ ** | 8.58 x $10^{-6}$ ** | 8.37 x $10^{-3}$ * | - | - | - |
| | Iself | [1, 3] | 0.25 | 0.27 | 0.18 | 0.03 * | 0.07 | 0.10 | 0.20 | 0.01 * | - | - |
| | Iothers | | 0.02 * | 8.58 x $10^{-6}$ ** | 8.58 x $10^{-6}$ ** | 8.58 x $10^{-6}$ ** | 5.72 x $10^{-5}$ ** | 5.72 x $10^{-5}$ ** | 8.58 x $10^{-6}$ ** | 8.58 x $10^{-6}$ ** | - | - |
| | Iself | [2, 3] | 0.18 | 0.26 | 0.06 | 0.02 * | 0.26 | 0.64 | 0.08 | 3.55 x $10^{-3}$ * | 0.08 | - |
| | Iothers | | 3.88 x $10^{-3}$ * | 0.30 | 0.30 | 2.26 x $10^{-5}$ ** | 7.54 x $10^{-3}$ * | 7.54 x $10^{-3}$ * | 1.61 x $10^{-5}$ ** | 8.58 x $10^{-6}$ ** | 0.73 | - |
| All vs 150 volumes | Iself | | 1.50 x $10^{-4}$ ** | 6.36 x $10^{-6}$ ** | 6.36 x $10^{-6}$ ** | 6.36 x $10^{-6}$ ** | 6.36 x $10^{-6}$ ** | 6.36 x $10^{-6}$ ** | 6.03 x $10^{-4}$ ** | 6.36 x $10^{-6}$ ** | 0.02 * | 0.73 |
| | Iothers | | 2.73 x $10^{-6}$ ** | 2.73 x $10^{-6}$ ** | 2.73 x $10^{-6}$ ** | 2.73 x $10^{-6}$ ** | 2.73 x $10^{-6}$ ** | 2.73 x $10^{-6}$ ** | 4.77 x $10^{-6}$ ** | 2.73 x $10^{-6}$ ** | 5.36 x $10^{-4}$ ** | 0.18 |

**Supplementary table S3. All volumes between-TR statistics summary table - *Iself and Iothers*.** For each comparison between TR combinations, *pFDR* values are reported for the all volumes analysis (*****: *pFDR* < 0.05, ******: *pFDR* < 0.001). Moreover, for each TR combination, *pFDR* values resulting from the comparison between the all volumes and the 150 volumes distributions are summarized. Wilcoxon signed rank was used for both self-identifiability (*Iself*) and others-identifiability (*Iothers*).

### S3. Whole-brain between-TR connectome fingerprinting: results on 150 volumes

PCA reconstruction on the single TR pairwise comparisons FC matrices revealed an optimal number of PCs close to the sample size (m* = 20.5 ± 1.1; **Supplementary table S4**, **Supplementary figure S1D**). Comparing observed *SR* and *Idiff* computed on the single identifiability matrices against their correspondent null distributions (Sareen et al., 2021) still resulted in a statistically significant effect (permutation testing, $p < 0.001$) for all TR combinations.

Moreover, the same trend as in the all volumes condition was found, according to which the higher the similarity between TRs, the higher the *Idiff*, *SR*, *Iself* and *Iothers* values (**Supplementary figure S5**, **Supplementary table S5, Supplementary table S6**).

| TR (s) | *m** | $R^2$ | $Idiff_{orig}$ | $Idiff_{recon}$ | $Iself_{recon}$ | $Iothers_{recon}$ | $SR_{orig}$ | $SR_{recon}$ |
|---|---|---|---|---|---|---|---|---|
| **[0.5, 0.7]** | 18 | 0.71 | 9.99 | 23.55 | 58.51 | 34.95 | 40.00 | 42.50 |
| **[0.5, 1]** | 20 | 0.77 | 8.76 | 21.05 | 57.50 | 36.45 | 47.50 | 47.50 |
| **[0.5, 2]** | 21 | 0.83 | 8.09 | 17.40 | 54.62 | 37.22 | 42.50 | 50.00 |
| **[0.5, 3]** | 21 | 0.82 | 7.35 | 15.50 | 50.78 | 35.31 | 32.50 | 42.50 |
| **[0.7, 1]** | 21 | 0.81 | 13.82 | 30.25 | 72.89 | 42.64 | 72.50 | 75.00 |
| **[0.7, 2]** | 21 | 0.85 | 14.34 | 27.62 | 72.75 | 45.13 | 82.50 | 85.00 |
| **[0.7, 3]** | 22 | 0.85 | 11.77 | 24.09 | 65.75 | 41.66 | 57.50 | 77.50 |
| **[1, 2]** | 20 | 0.84 | 17.04 | 32.32 | 81.91 | 49.59 | 75.00 | 85.00 |
| **[1, 3]** | 21 | 0.83 | 16.33 | 32.79 | 78.59 | 45.80 | 82.50 | 90.00 |
| **[2, 3]** | 20 | 0.85 | 19.88 | 36.77 | 85.95 | 49.18 | 82.50 | 90.00 |

**Supplementary table S4. 150 volumes between-TR analysis summary table.** For each TR combination, values of the percentage differential identifiability (*Idiff*), self-identifiability (*Iself*), others-identifiability (*Iothers*) and success rate (*SR*) are reported for the 150 volumes condition. *Orig* values were computed on the identifiability matrices derived from the original (before Principal Component Analysis reconstruction) functional connectivity (FC) matrices, while *Recon* values were extracted from the FC matrices reconstructed by using the optimal number of principal components (m*), for which explained variance ($R^2$) is also reported.

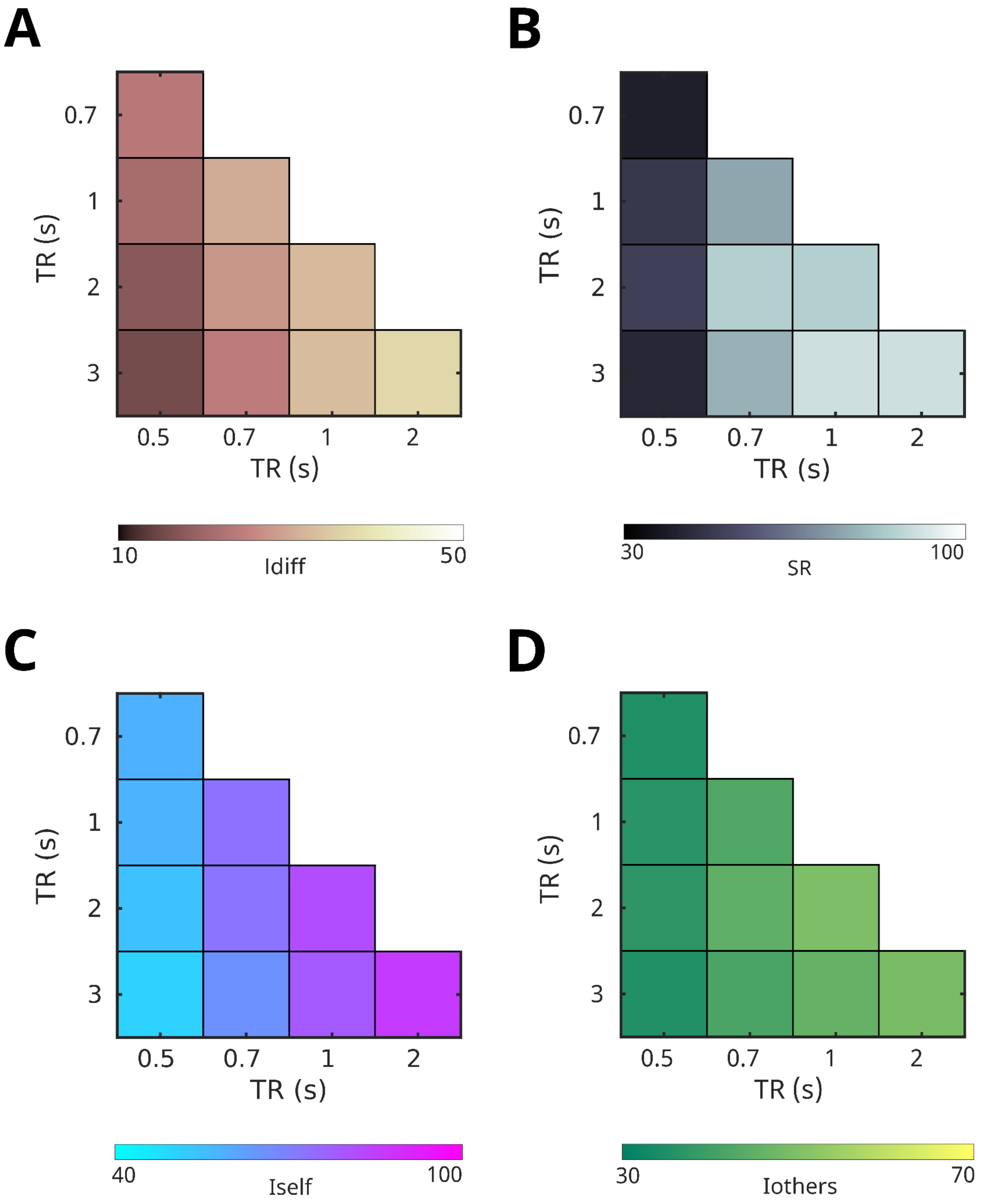


**Supplementary figure S5. 150 volumes between-TR fingerprinting.** Confusion matrices of the *Idiff* (**A**), *SR* (**B**), *Iself* (**C**) *and Iothers* (**D**) values computed from the between-TR PCA-reconstructed identifiability matrices (**Supplementary figure 3B**).

| | [TR*i*, TR*j*] (s) | [0.5, 0.7] | [0.5, 1] | [0.5, 2] | [0.5, 3] | [0.7, 1] | [0.7, 2] | [0.7, 3] | [1, 2] | [1, 3] |
|---|---|---|---|---|---|---|---|---|---|---|
| Idiff | **[0.5, 1]** | 0.50 | - | - | - | - | - | - | - | - |
| SR | | 0.65 | - | - | - | - | - | - | - | - |
| Idiff | **[0.5, 2]** | 0.06 | 0.31 | - | - | - | - | - | - | - |
| SR | | 0.97 | 1.00 | - | - | - | - | - | - | - |
| Idiff | **[0.5, 3]** | 0.07 | 0.24 | 0.48 | - | - | - | - | - | - |
| SR | | 1.00 | 0.97 | 1.00 | - | - | - | - | - | - |
| Idiff | **[0.7, 1]** | 0.20 | 0.08 | **0.01 *** | **0.01 *** | - | - | - | - | - |
| SR | | **0.01 *** | 0.12 | 0.08 | **0.01 *** | - | - | - | - | - |
| Idiff | **[0.7, 2]** | 0.47 | 0.26 | **0.03 *** | **0.02 *** | 0.32 | - | - | - | - |
| SR | | **0.01 *** | **0.05 *** | **0.05 *** | **0.02 *** | 1.00 | - | - | - | - |
| Idiff | **[0.7, 3]** | 0.80 | 0.44 | 0.06 | 0.06 | 0.05 | 0.17 | - | - | - |
| SR | | **0.01 *** | 0.08 | **0.03 *** | **0.03 *** | 1.00 | 1.00 | - | - | - |
| Idiff | **[1, 2]** | 0.19 | 0.07 | **0.01 *** | **0.01 *** | 0.80 | 0.24 | **0.03 *** | - | - |
| SR | | **0.01 *** | **0.05 *** | **0.02 *** | **0.01 *** | 0.94 | 1.00 | 1.00 | - | - |
| Idiff | **[1, 3]** | 0.14 | **0.04 *** | **0.01 *** | **0.01 *** | 0.90 | 0.26 | **0.04 *** | 0.72 | - |
| SR | | **0.01 *** | **0.03 *** | **0.01 *** | **0.01 *** | 0.94 | 1.00 | 1.00 | 1.00 | - |
| Idiff | **[2, 3]** | 0.06 | **0.01 *** | **$3 \times 10^{-3}$ *** | **0.01 *** | 0.24 | **0.02 *** | **0.01 *** | 0.08 | 0.07 |
| SR | | **0.01 *** | **0.02 *** | **0.01 *** | **0.01 *** | 0.65 | 0.94 | 0.83 | 1.00 | 1.00 |

**Supplementary table S5. 150 volumes between-TR statistics summary table - *Idiff* and *SR*.** For each comparison between TR combinations, pFDR values are reported for the 150 volumes analysis (*: pFDR < 0.05, **: pFDR < 0.001). Wilcoxon signed rank was used for differential identifiability (*Idiff*), while McNemar's test for proportions in paired samples was employed for success rate (*SR*). Statistics refer to functional connectivity matrices reconstructed by using the optimal number of principal components.

| | [TR*i*, TR*j*] (s) | [0.5, 0.7] | [0.5, 1] | [0.5, 2] | [0.5, 3] | [0.7, 1] | [0.7, 2] | [0.7, 3] | [1, 2] | [1, 3] |
|---|---|---|---|---|---|---|---|---|---|---|
| Iself | [0.5, 1] | 0.72 | - | - | - | - | - | - | - | - |
| Iothers | | 0.08 | - | - | - | - | - | - | - | - |
| Iself | [0.5, 2] | 0.20 | 0.50 | - | - | - | - | - | - | - |
| Iothers | | 0.10 | 0.15 | - | - | - | - | - | - | - |
| Iself | [0.5, 3] | 0.08 | 0.12 | 0.13 | - | - | - | - | - | - |
| Iothers | | 0.67 | 0.08 | 3.59 x $10^{-3}$ * | - | - | - | - | - | - |
| Iself | [0.7, 1] | 0.02 * | 0.02 * | 3.80 x $10^{-3}$ * | 1.55 x $10^{-3}$ * | - | - | - | - | - |
| Iothers | | 1.01 x $10^{-5}$ ** | 1.63 x $10^{-4}$ ** | 4.05 x $10^{-3}$ * | 8.58 x $10^{-5}$ ** | - | - | - | - | - |
| Iself | [0.7, 2] | 0.02 * | 0.01 * | 1.99 x $10^{-3}$ * | 1.11 x $10^{-3}$ * | 0.90 | - | - | - | - |
| Iothers | | 1.01 x $10^{-5}$ ** | 2.15 x $10^{-5}$ ** | 7.80 x $10^{-6}$ ** | 7.80 x $10^{-6}$ ** | 1.39 x $10^{-3}$ * | - | - | - | - |
| Iself | [0.7, 3] | 0.15 | 0.12 | 0.01 * | 4.26 x $10^{-3}$ * | 0.02 * | 0.01 * | - | - | - |
| Iothers | | 1.32 x $10^{-4}$ ** | 8.23 x $10^{-4}$ ** | 6.98 x $10^{-3}$ * | 7.80 x $10^{-6}$ ** | 0.19 | 5.22 x $10^{-5}$ ** | - | - | - |
| Iself | [1, 2] | 1.11 x $10^{-3}$ * | 1.11 x $10^{-3}$ * | 2.32 x $10^{-4}$ ** | 2.15 x $10^{-4}$ ** | 0.02 * | 0.02 * | 2.21 x $10^{-4}$ ** | - | - |
| Iothers | | 7.80 x $10^{-6}$ ** | 7.80 x $10^{-6}$ ** | 7.80 x $10^{-6}$ ** | 7.80 x $10^{-6}$ ** | 1.01 x $10^{-5}$ ** | 1.32 x $10^{-4}$ ** | 7.80 x $10^{-6}$ ** | - | - |
| Iself | [1, 3] | 4.54 x $10^{-3}$ * | 3.20 x $10^{-3}$ * | 3.14 x $10^{-4}$ ** | 2.32 x $10^{-4}$ ** | 0.46 | 0.27 | 1.99 x $10^{-3}$ * | 0.12 | - |
| Iothers | | 1.01 x $10^{-5}$ ** | 2.86 x $10^{-5}$ ** | 5.22 x $10^{-5}$ ** | 2.15 x $10^{-5}$ ** | 0.01 * | 0.58 | 8.58 x $10^{-5}$ ** | 2.15 x $10^{-5}$ ** | - |
| Iself | [2, 3] | 2.68 x $10^{-4}$ ** | 6.00 x $10^{-4}$ ** | 2.15 x $10^{-4}$ ** | 2.32 x $10^{-4}$ ** | 3.58 x $10^{-3}$ * | 3.80 x $10^{-3}$ * | 2.21 x $10^{-4}$ ** | 0.03 * | 4.54 x $10^{-3}$ * |
| Iothers | | 1.01 x $10^{-5}$ ** | 7.80 x $10^{-6}$ ** | 1.01 x $10^{-5}$ ** | 7.80 x $10^{-6}$ ** | 1.09 x $10^{-4}$ ** | 8.23 x $10^{-4}$ ** | 7.80 x $10^{-6}$ ** | 0.52 | 4.84 x $10^{-4}$ ** |

**Supplementary table S6. 150 volumes between-TR statistics summary table - *Iself* and *Iothers*.** For each comparison between TR combinations, *pFDR* values are reported for the 150 volumes analysis (*****: *pFDR* <

0.05, **: *pFDR* < 0.001). Wilcoxon signed rank was used for both self-identifiability (*Iself*) and others-identifiability (*Iothers*).

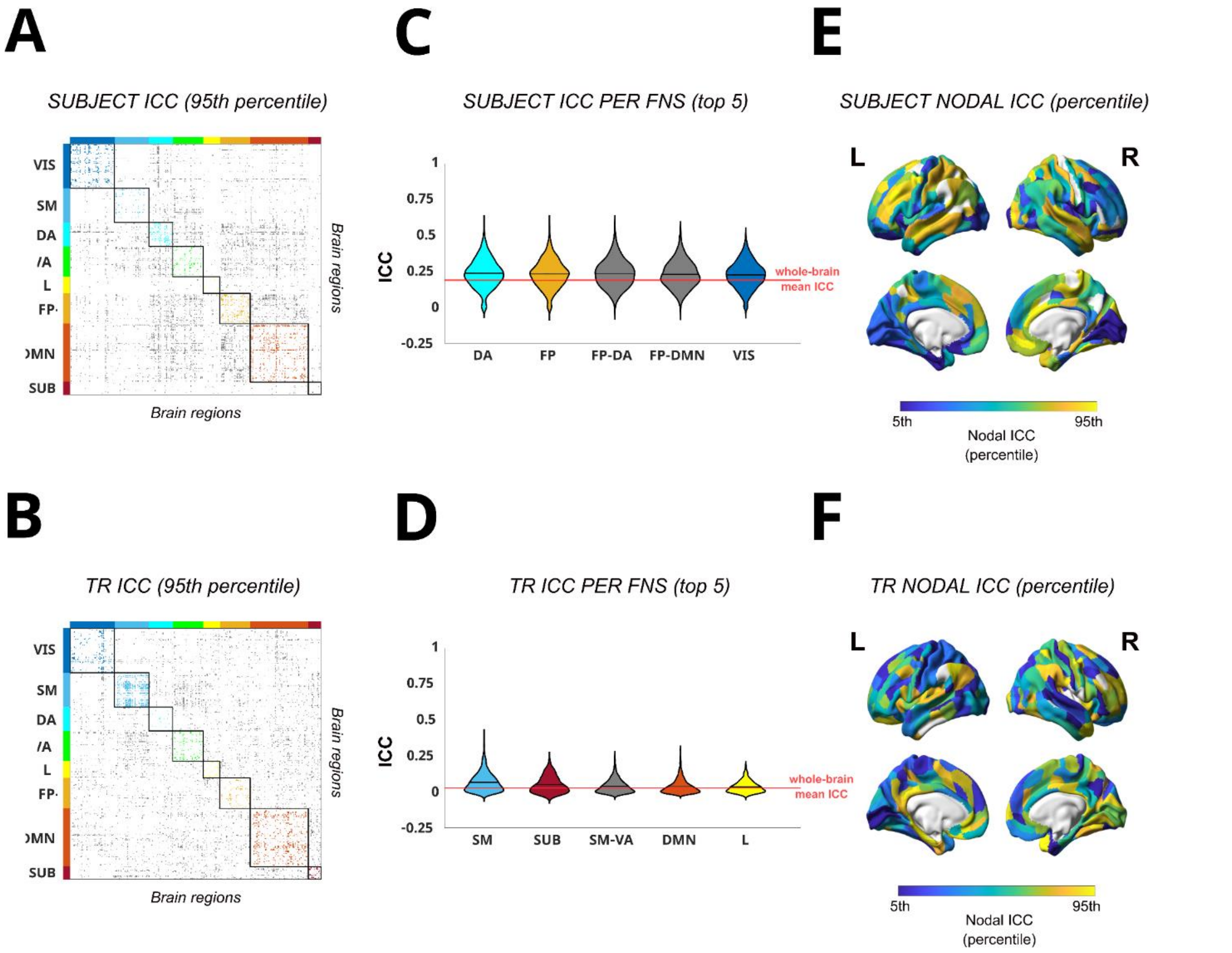


**Supplementary figure S6. 150 volumes edgewise Intra-class correlation (ICC) analysis of subject identifiability and task identifiability.** (**A-B**) Edgewise subject (**A**) and TR ICC (**B**) matrices, showing only functional connections with ICC values significantly higher than the mean distribution (that is, lying in the 95th percentile). The brain regions are ordered according to Yeo's (Yeo et al., 2011) functional resting state networks (FNs): Visual (VIS), Somato-Motor (SM), Dorsal Attention (DA), Ventral Attention (VA), Limbic system (L), Fronto-Parietal (FP), Default Mode Network (DMN), and subcortical regions (SUB). The colored dots refer to within FNs networks edges, while gray dots refer to between FNs networks edges, as in Amico & Goñi (2018). (**C-D**) Violin plots of edgewise subject (**C**) and TR (**D**) ICC distributions for the 5 FNs with the highest mean ICC value. Each colored violin plot indicates a different within FN, while gray violin plots indicate between FNs ICC distributions. The horizontal solid black line within each violin plot indicates the mean value of each distribution; the solid red line across the violin plots, instead, indicates the whole-brain mean ICC value, as in Amico & Goñi (2018). The most prominent FNs for subject's identifiability (**C**) resulted: DA, FP, VIS, and the FP-DA, FP-DMN interactions. For TR identifiability, the most relevant FNs were (**D**): SM, SUB, L, DMN, and SM-VA interactions. (**E-F**) Brain render of nodal ICC, computed as the column-wise mean of the edgewise ICC matrices for both subject (**E**) and TR (**F**) ICC, and represented at 5th - 95th percentile threshold. Nodal ICC gives an assessment of the overall prominence of each brain region for subject's and TR identifiability. All plots refer to the 150 volumes analysis.

### S4. Whole-brain within-TR connectome fingerprinting: the effect of band-pass filtering

During data pre-processing, a standard temporal band-pass filtering ([0.01-0.3] Hz) was applied regardless of the acquisitions' temporal resolution. At TR 0.3 s, however, this band-pass filter covers the whole frequency range of the original spectrum and its aliasing introduced by the digital sampling (see **Appendix**). This does not happen for the remaining TRs taken into account in our work. To exclude the possibility that this aspect may have introduced a bias in the data which have ultimately led to a statistically significant higher *Idiff* at TR 3 s in comparison to TR 0.7, 1 and 2 s, we rerun the analysis after applying a [0.01-0.16] Hz band-pass filter during the preprocessing of all the acquisitions' data. The value 0.16 Hz corresponds to the Nyquist frequency at TR 3 s. We found no qualitatively different effects between the two differently band-pass filtered data, so we conclude that identifiability effects of the temporal resolution in the within-TR fingerprinting analysis were due signal sampling (see **Appendix**) rather than temporal filter effects.

### Appendix

In the current study, we employed the brain functional connectome fingerprinting framework to investigate how the temporal resolution at which resting-state functional MRI (rs-fMRI) data were acquired affected the possibility to uniquely identify one individual among a sample of 20 healthy participants only on the basis of their intrinsic functional connectivity. Each volunteer underwent five whole-brain rs-fMRI acquisitions with varying TR (0.5 s, 0.7 s, 1 s, 2 s, and 3 s). After preprocessing, the differential identifiability framework introduced by Amico and Goñi (2018) was applied to investigate test-retest reliability within the two halves of each TR acquisition separately. Despite the high identifiability achieved regardless of the temporal resolution at which data were acquired, fingerprinting analysis was still sensitive to the fMRI acquisition protocol, and in particular to TR. We found (**Figure 2B**) that *diff* at TR 0.5 s was significantly higher than *Idiff* from all other TRs, and that TR 3 s significantly outperformed TR 0.7, 1 and 2 s in *Idiff*. According to our interpretation, the observed TR-dependent effects are due to the sampling of the physiological noise, which is aliased into the lower frequency range, and added to the neural signal. This results in a series of overlapping frequencies whose source identity cannot be reconstructed back. In other words, an increase in the signal-to-noise ratio is observed, and, thus, in the loss of information which could potentially contribute to distinguishing one individual’s functional connectivity profile from another’s.

To better explain our interpretation, let us assume a simple signal measurement model where the continuous MRI signal is sampled at every TR. This can be modeled as an impulse train with a rectangular window, having a width equal to the sampling period, that is [0–TR]. The power spectral density of this impulse (a squared *sinc* function) can be used to evaluate the frequencies that contribute to the sampled signal. **Figure A1** below depicts the spectral density in frequency domain for each of the TRs considered: the black line corresponds to the analog signal, while the dashed colored lines represent the positive (magenta) and negative (red) sidebands introduced by the sampling impulse. The gray area of the original spectrum gets aliased to the baseband by the sidebands contributions, i.e., the magenta and red shaded areas. This shows that the largest aliasing from the original spectrum to the sampled signal originates from the spectral density ranging from the Nyquist frequency, that is half of the sampling rate, and the sampling rate itself, that is the TR expressed as frequency.

As it emerges from the simulation, at both TR 0.5 s and TR 3 s, the spectral density of the aliasing ([1–2] Hz and [0.17–0.33] Hz, respectively) avoids the range [0.33–1] Hz, which contains most of the physiological contributions, with respiratory brain pulsations lying respectively in [0.25–0.35] Hz frequency band, and cardiac noise lying in and [0.8–1.1] Hz frequency band. As a consequence, we observe that the participants' functional connectivity profiles are maximally identifiable at TR 0.5 s and TR 3 s, with lower identifiability for intermediate TRs.

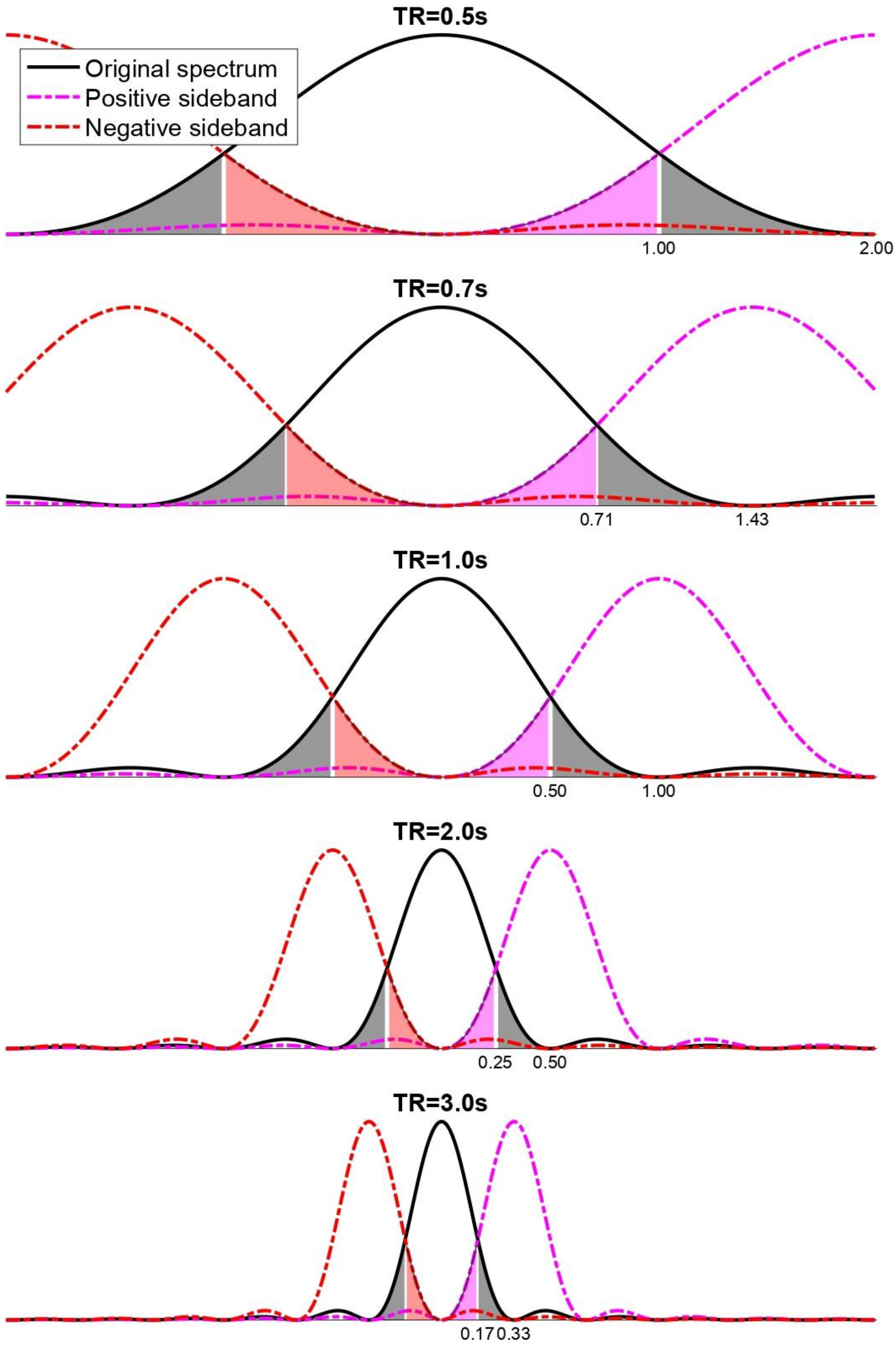


**Figure A1. Power spectrum frequency response of MRI signal sampled at different TRs.** Spectral responses in frequency space (Hz) of different impulse [0-TR] signals. A baseline response (continuous black line) includes positive (magenta dashed line) and negative (red dashed line) sidebands introduced due to sampling. The gray area of the original spectrum gets aliased to the baseband by the sideband contributions (red and magenta shaded areas), with the largest aliasing originating in the frequency range [1/(2TR),1/TR], shown on each plot.
